# Spatially resolved single-cell atlas of the lung in fatal Covid19 in an African population reveals a distinct cellular signature and an interferon gamma dominated response

**DOI:** 10.1101/2023.11.16.566964

**Authors:** James Nyirenda, Olympia Hardy, João Da Silva Filho, Vanessa Herder, Charalampos Attipa, Charles Ndovi, Memory Siwombo, Takondwa Namalima, Leticia Suwedi, Watipenge Nyasulu, Thokozile Ngulube, Deborah Nyirenda, Leonard Mvaya, Joseph Phiri, Dennis Chasweka, Chisomo Eneya, Chikondi Makwinja, Chisomo Phiri, Frank Ziwoya, Abel Tembo, Kingsley Makwangwala, Stanley Khoswe, Peter Banda, Ben Morton, Orla Hilton, Sarah Lawrence, Monique Freire dos Reis, Gisely Cardoso Melo, Marcus Vinicius Guimaraes de Lacerda, Fabio Trindade Maranhão Costa, Wuelton Marcelo Monteiro, Luiz Carlos de Lima Ferreira, Carla Johnson, Dagmara McGuinness, Kondwani Jambo, Michael Haley, Benjamin Kumwenda, Massimo Palmarini, Kayla G. Barnes, Donna M. Denno, Wieger Voskuijl, Steve Kamiza, Kevin Couper, Matthias Marti, Thomas Otto, Christopher A. Moxon

## Abstract

Postmortem single-cell studies have transformed understanding of lower respiratory tract diseases (LRTD) including Covid19 but there is almost no data from African settings where HIV, malaria and other environmental exposures may affect disease pathobiology and treatment targets. We used histology and high-dimensional imaging to characterise fatal lung disease in Malawian adults with (n=9) and without (n=7) Covid19, and generated single-cell transcriptomics data from lung, blood and nasal cells. Data integration with other cohorts showed a conserved Covid19 histopathological signature, driven by contrasting immune and inflammatory mechanisms: in the Malawi cohort, by response to interferon-gamma (IFN-γ) in lung-resident alveolar macrophages, in USA, European and Asian cohorts by type I/III interferon responses, particularly in blood-derived monocytes. HIV status had minimal impact on histology or immunopathology. Our study provides data resources and highlights the importance of studying the cellular mechanisms of disease in underrepresented populations, indicating shared and distinct targets for treatment.

## Introduction

There has been significant progress towards creation of a human cell atlas utilising scRNA-sequencing (scRNA-seq) and high-dimensional cellular imaging data^1,2^. The human cell atlas is transforming our understanding of cells and their states in health and disease and is rapidly becoming a major resource for the development of novel treatments and vaccines^3^. Yet, data within this atlas is heavily biased towards populations in the Northern hemisphere. Populations in sub–Saharan Africa (SSA) are particularly underrepresented^4^. Genetic and environmental factors may lead to important differences in cell development and cell-compositions in different organs, thus effecting cellular responses to diseases, vaccines and therapies^5,6^. Capturing data from SSA populations is critical to assure that everyone can benefit from the treatment advances derived from the human cell atlas.

Immunomodulation plays a critical role in Covid19 outcomes. Single-cell data from lung tissue facilitated identification of specific immunomodulatory targets^3,7-12^. Apart from our high-dimensional imaging study from a Brazilian cohort^13^ these data are, thus far, exclusively from populations in Northern hemisphere, similar to most clinical trial data validating their efficacy. For future waves or epidemics of SARS-CoV2 or related viruses, this knowledge gap needs to be addressed. Indeed, given minimal intensive care, the benefit of preventing progression to or deterioration from severe disease by immunomodulation is even more important in SSA. While immunomodulatory therapies can be lifesaving, they can also be harmful^14^. Immunomodulation has focused on two opposing strategies: augmenting the inflammatory response to aid viral clearance or attenuating inflammatory response to reduce pathogenic hyperinflammation. Extensive studies in northern hemisphere cohorts have established that, by the time patients present with life threatening illness, viral loads are declining, hyperinflammation generally predominates and thus anti-inflammatory interventions are more effective^14,15^. Given evidence that repeated exposure to malaria and other parasitic infections can induce immune tolerance^16-18^, we hypothesised that the balance may be different in patients in SSA where these infections are more prevalent. While sometimes this clinical context may be protective, in those who progress to severe disease, a tolerance-skewed response might blunt immune-mediated viral clearance, leading to a more viral-driven pathology. However, the reverse is also possible. High pathogen exposure can induce an accelerated inflammatory response on re-exposure to pathogens^5^. Either scenario might impact cellular responses driving pathogenesis in the lung and have important implications for informing which therapy may be effective in SSA populations.

To address some of these knowledge gaps we conducted an autopsy study in well-characterised patients at a large public hospital in Malawi, a low-income country in SSA with high rates of malaria, TB and HIV. We undertook detailed histopathological analysis and scRNA-seq on lung, blood and nasal cells and imaging mass cytometry (IMC) to spatially resolve the immune landscape of the lung (Fig.1a). We conducted all tissue processing, cell dissociation and scRNA-seq library preparation on site in Malawi, with much of the data prepared on fresh samples. There are so far no studies from any settings that included characterisation across all these modalities. Thus, to fully understand the context of our data in contrast to other populations, we needed to use data from patient cohorts from different regions of the world to enable comparisons (Fig.1b). Taken together, our data highlight how Covid19 has a similar histopathological pattern in our SSA cohort to other Northern and Southern Hemisphere cohorts. However, we found a contrasting immune response signature in the SSA cohort, driven by proliferation of lung-resident alveolar macrophages and interferon gamma (IFN-γ).

**Figure 1.**
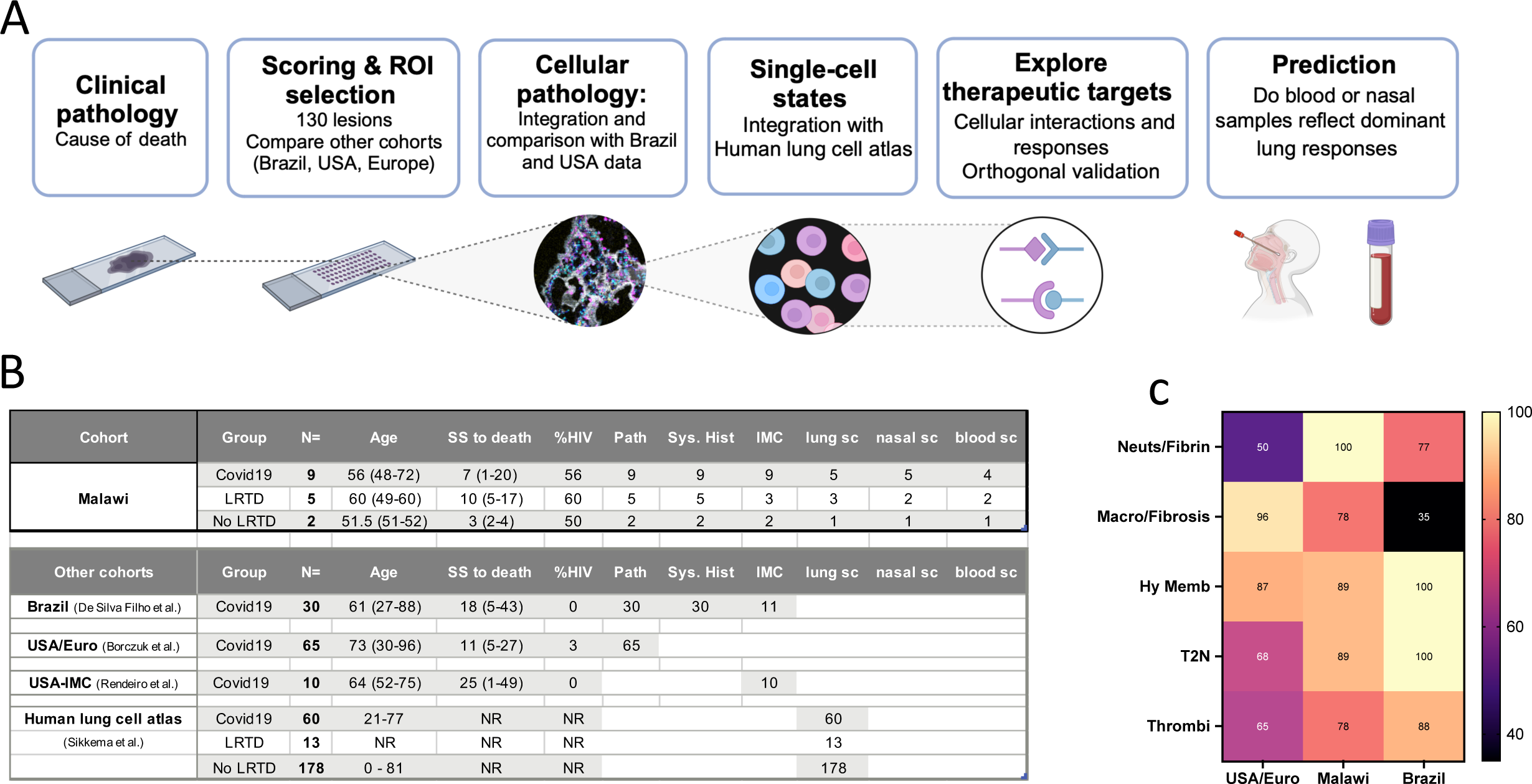
Study overview, overview of our cohort and comparator cohorts and histological lesion comparison with other cohorts. **A)** Overview of study approach, image created using BioRender; **B)** Summary of the characteristics of our Malawi cohort versus published cohorts that we have used for different comparisons. **C)** Heatmap shows the proportion of cases in the three cohorts (USA/European; Malawi; Brazil) that have each given lesion type. Abbreviations: SS to death = symptom start to death in days. Path = Pathology. indicates the number of cases included in each cohort in which postmortem pathological features are described. Sys. Hist. = Systematic histopathology, denotes the number of cases included in each cohort with scoring of the frequency and severity of different lesions scored based on pre-defined criteria. IMC = Imaging Mass Cytometry, number of cases with data for this. Lung sc = lung cell single-cell RNA-seq, denotes the number of cases with scRNA-seq data from lung tissue. nasal sc = Nasal cell single-cell RNA-seq, denotes the number of cases with scRNA-seq data from nasal tissue. blood sc = blood cell single-cell RNA-seq, number of cases with this data

## Results

### Clinical and histopathological analysis identifies a conserved histopathological signature of Covid19 in patients from SSA, in agreement with other Covid19 cohorts

We recruited patients with fatal illness aged 45-75 admitted to Queen Elizabeth Central Hospital, Blantyre (October 2020 and July 2021) and stratified into three groups based on clinical characteristics: 1) Covid19 acute respiratory distress syndrome (ARDS) (n=9), 2) lower respiratory tract disease (LRTD) (n=5) with ARDS of diverse non-Covid19 aetiology, 3) non-LRTD (n=2) (Fig.1b, Extended Data Table 1, see methods). Like other cohorts, fatal Covid19 cases were generally overweight or obese (78%) and several cases had type 2 diabetes (44%), whereas LRTD and non-LRTD cases were generally underweight. HIV was common across all groups. No new diagnoses of HIV were made, and all infected cases had been on highly active anti-retroviral treatment until stopping treatment during the pandemic, reflecting challenges with access to healthcare and in keeping with low CD4 counts (median 134cells/mm^3^).

Autopsies were conducted through minimally invasive tissue sampling (MITs)^19^ using wide-bore needle-biopsies. Lung, liver and brain samples were obtained in all 16 cases, bone marrow in 15 cases and spleen in 8 cases. A pathologist (S.K.) read haematoxylin and eosin-stained tissue slides alongside patients’ history and antemortem lab results, per standard clinical practice. In the lung in the Malawi Covid19 cohort the pathologist identified classical features of Covid19 described in other cohorts^20-27^, which were absent or less frequent in LRTD cases (Supplemental Data Fig.1). In contrast, in other organs there were no Covid19 specific changes (Supplemental Data Table 1), focusing our further investigations on the lung. Then, two additional pathologists (V.H., C.A.), blinded to diagnosis, scored the lung pathology in all sixteen patients using more detailed semi-quantitative criteria developed by us previously^13^. Within our Covid19 cases, type II pneumocyte hyperplasia, vascular congestion, syncytia, granulation of tissue and lymphocyte infiltration were all significantly more common and severe than in the non-Covid19 LRTD group; in contrast neutrophils were more numerous in LRTD cases (Extended Data Fig.1a). There was no significant difference in histopathology due to HIV status (Extended data Fig.1b).

Unfortunately, a lack of international consensus criteria for assessing Covid19 lung pathology, and of studies with systematic scoring, prevented quantitative comparison with other cohorts to assess similarities and differences. Therefore, we compared proportions of different pulmonary lesion types with a study that combined cohorts from Europe and the USA^21^, and with our published Brazilian cohort^13^ (Fig.1c). Acute alveolar changes, defined by neutrophil infiltration and fibrin-deposition were more frequent in the Malawi and Brazil cohorts than the USA cohort. “Chronic” alveolar changes with monocytes, macrophages or fibrosis were detected more frequently in the USA and Malawi cohort. In the Malawi cohort, “chronic” disease was predominantly characterized by macrophage and monocytes and there was less fibrosis than in the USA and Brazil cohorts. The detection of macrophages by histology in a high proportion of Malawi cases, despite a very short duration of illness, fits with studies that observed inflammatory monocyte/macrophage populations in fatal Covid19 cases with a short durations of illness^8,28,29^, indicating a departure from classical “acute” to “chronic” changes. Thus, despite a short duration from illness to death, and demographic differences, cases in our Malawi cohort exhibited classical Covid19 lung pathology.

### High-plex imaging highlights that Covid19 immunopathology is associated with alveolar macrophages whereas LRTD is associated with neutrophils

To give cellular context to the histopathological features we made tissue microarrays, identifying 130 representative regions of interest containing specific pathological lesions or normal lung areas (9 Covid, 3 LRTD, 2 Non-LRTD cases). Tissue samples were analysed by IMC using a 39 metal-conjugated antibody panel (Supplemental information) that we previously optimised for staining in a Brazil cohort^13^, an anti-SARS-CoV2 Spike (SARS-CoV2-S) antibody validated in lung tissue^13,30^. After cell-segmentation and quality control, we annotated 76,369 cells. Cells were divided into major subtypes and then subtypes classified based on markers of activation, differentiation, proliferation and apoptosis (Fig.2a, Extended data Fig.2a).

**Figure 2.**
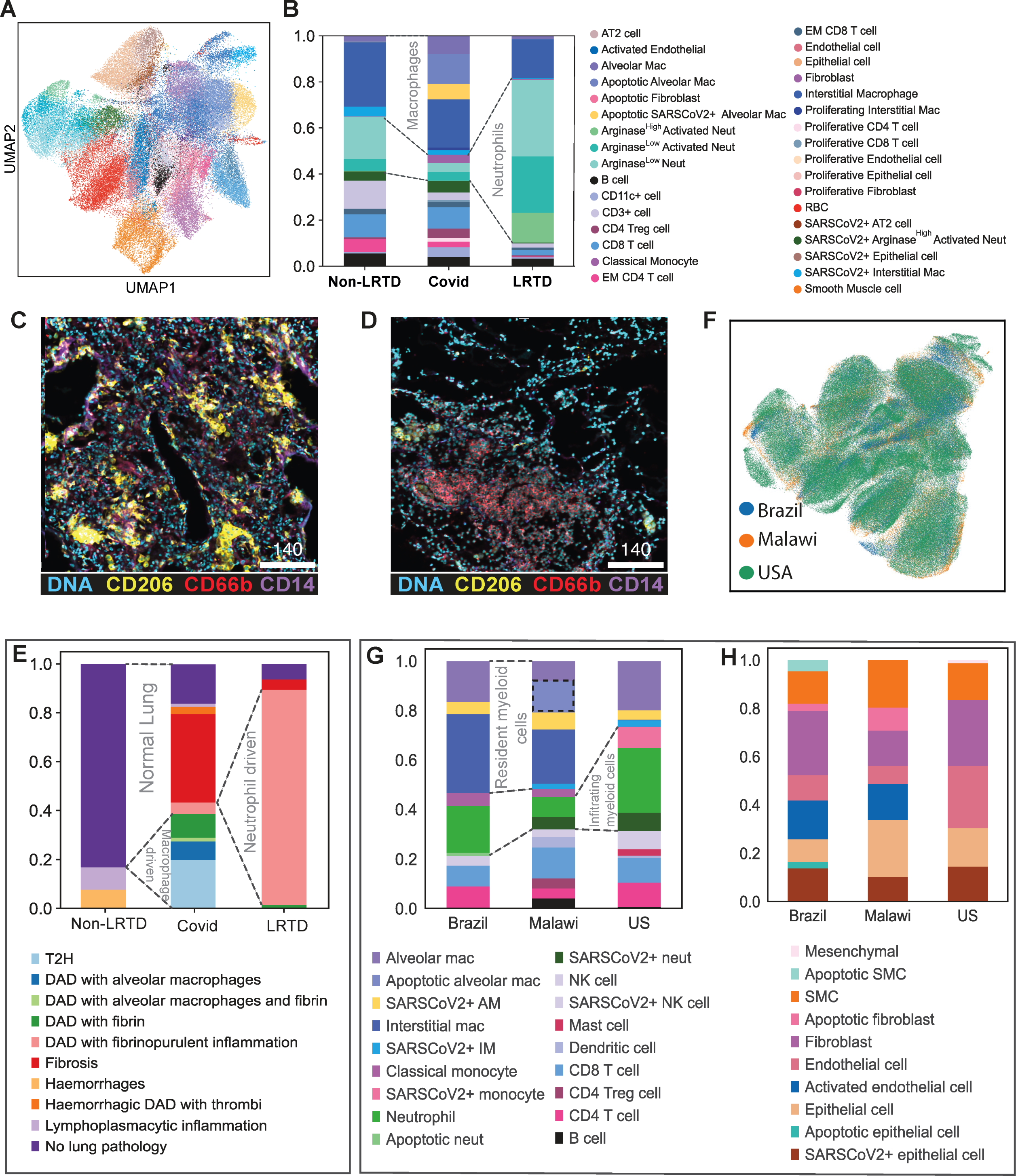
Imaging Mass Cytometry (IMC) reveals an immunopathological landscape of Covid19 in Malawi cases driven by alveolar macrophages A) UMAP embedding of the cell types identified in the lung samples by IMC, after supervised assignment to major cell types. Each major cell type was clustered and resulting clusters were annotated and merged to extract the final set of cell types. Colour key for cell types is on right hand side of B. **B)** Frequency of the immune cell types identified in the post-mortem lung samples by IMC according to clinical groups. The stacked bar plot shows the averaged frequency of the cell types by grouping the values from regions of interest (ROIs) according to the clinical groups. Dashed lines highlight principle differences in major cell populations between COVID-19 and other respiratory disease groups. **C)** Representative denoised IMC images from COVID19 case shows high levels of CD206^+^ macrophages (yellow) and lower levels of neutrophils (CD66b, red) and monocytes (CD14, purple). **D)** Representative denoised IMC images from LRTD case shows high levels of high levels of neutrophils (CD66b, red) and low levels of CD206^+^ macrophages (yellow). **E)** Frequency of histopathological lesions based on matched H&E and IMC analysis of post-mortem lung samples from the different clinical groups. The cellular composition and frequency of different cell types is indicated in Extended data Fig. 3F. **F)** UMAP embedding of the integrated IMC lung data from the Brazil, USA and Malawi COVID-19 cohorts. **G)** Comparison of immune cell frequencies in IMC data from Brazil, Malawi and USA cohorts after integration shown in F, some major cell group differences are highlighted by dotted lines. Dashed box highlights apopototic alveolar macrophages which are only present in the Malawi cohort. **H)** Comparison of stromal cell frequencies in IMC data from Brazil, Malawi and USA cohorts after integration shown in F.

In our Malawi cohort, neutrophils (CD66b^pos^CD11b^pos^CD14^neg/low^) were significantly more numerous in the LRTD cases (49.6%) than in the non-LRTD (21.1%) or Covid19 cases (16.1%, adjusted p=<0.001) (Fig.2b,d, Extended Data Table2 and Fig.2e). Reciprocally, macrophages were increased in Covid19 (44.1%) compared to LRTD (30.4%) and non-LRTD cases (23.6%; adjusted p=<0.0001, Fig.2b,c, Extended Data Table 2). In contrast to data in prior published USA and European cohorts^28,30^, these were predominantly alveolar macrophages (CD206^high^CD163^high^Iba1^low^MHCII^low^CD14^neg^) with a lower number of monocyte derived CD14^high/int^ cells.

There were no consistent differences in T-cell numbers between the Covid19, LRTD and non-LRTD disease groups, but among Covid19 cases there was an expansion in Tregs and proliferating T-cells and a decrease in the ratio of effector memory (CD45RO^high^) to naïve (CD45RO^low^) CD8 T-cells (Fig.2b, Extended data Table 2). B-cell numbers were not markedly different in Covid19, although we had few B-cell markers (Fig.2b). Consistent with vascular pathology visualised by histology (fibrin deposition and thrombosis), there was increased endothelial cell activation in Covid19 compared with LRTD and non-LRTD cases (Fig.2b, Extended data Table2). Alveolar macrophages were the most common SARS-CoV2-S+ immune cell, followed by Arg^high^ neutrophils and interstitial macrophages (Fig.2b, Extended data Table 2). In the stromal compartment type 2 pneumocytes (AT2) and epithelial cells were the most frequent SARS-CoV2-S+ cells. We found no SARS-CoV2-S+ endothelial cells or fibroblasts. Surprisingly, total numbers of SARS-CoV2-S+ cells were lower in HIV+ cases (Extended data Fig.3c and Table 2).

Exploiting the spatial resolution of IMC and regions of interest selected based on a dominant pathological lesion, we characterised cellular compositions of lesion-types (Extended data Fig.2f), then quantified lesion-type levels by group (Fig.2e). Type II pneumocyte hyperplasia was specific to the Covid19 group. Diffuse alveolar damage occurred in both LRTD and Covid19 but had different compositions indicating different pathological processes. In LRTD, diffuse alveolar damage was mainly associated with neutrophil-driven fibrinopurulent inflammation; in Covid19, it had a more heterogenous immune-cell composition, dominated by the presence of macrophages, except fibrin-containing lesions which were neutrophilic. Together these implicate macrophages in alveolar damage and lung parenchymal processes and neutrophils in coagulopathic processes.

### Integration of high-plex data across Covid19 cohorts highlights shared different myeloid compositions and low levels of cells containing SARS-CoV2 antigen in Malawi cases

To systematically compare how the cell composition, degree of inflammation and amount of virus differed between our Malawi cohort and other cohorts we integrated data from Covid19 cases in our cohort (n=9) with two other cohorts that used IMC (Fig.2f): our Brazil cohort (n=11) that employed the same antibody panel^22^ and a USA cohort^30^ (n=10) that used several of the same markers including the same SARS-CoV2-S antibody. There were many similarities in cell proportions between the three cohorts but also significant differences (Fig2g,h). In the myeloid compartment there was a predominance of tissue-resident myeloid cells in the Malawi cohort versus infiltrating cells in the USA cohort, the Brazil cohort showed an intermediate phenotype. The USA cohort had the highest proportion of neutrophils (26.28%; adjusted p=2.64x10^-12^), and Malawi cohort the fewest (7.9%, Brazil 19%). Monocyte/macrophages were more frequent in the Brazil (58.5%) and Malawi (55.1%) than USA cohort (35.1%). Notably, the Malawi cohort had a high proportion of apoptotic alveolar macrophages (12.9%) which were absent in the other two cohorts, indicating that there may be specific macrophage responses in the Malawi cohort. Endothelial activation was prominent in the Brazil and Malawi cohorts, and virtually absent in the USA cohort (Fig.2h). In the stromal compartments there was a lower proportion of fibroblasts in the Malawi cohort, in keeping with lower levels of fibrosis on histology.

SARS-CoV-2-S antigen gives an indication of the quantity of viral material, although does not distinguish cells with replicating virus. We hypothesised that a tolerance-skewed immune response might lead to higher levels of virus in Malawi cases but did not find evidence of this. The USA cohort had the highest number of SARS-CoV2+ immune cells (25.7%, Malawi 13.9, Brazil 5%), These were principally monocytes and neutrophils in the USA cohort versus alveolar and interstitial macrophages in the Brazil and Malawi cohorts (Fig.2g, Extended Data Table 4). In the stromal compartment SARS-CoV2-S was detected in epithelial cells in all three cohorts but were significantly lower in the Malawi (Fig.2h)(6.3%, p=1.18x10^-10^) compared to Brazil (13.1%) and USA (12.1%) cohorts.

We found no evidence of blunted immune response or higher viral loads in the Malawi versus Brazil and USA cohort. The immune response signatures identified resident macrophages in the Malawi cohort versus neutrophil and monocyte infiltration in the USA cohort. A distinct apoptotic alveolar macrophage population in the Malawi cohort, together with the prominence of alveolar macrophages in lung lesions (Fig.2e), led us to consider whether these cells have differences in their inflammatory response in the Malawi cohort.

### Pulmonary cell scRNA-seq reveals low levels of viral RNA and an IFN-γ dominated response in the Malawi cohort

To explore cellular responses in the lung at greater depth in our Malawi cohort, including in alveolar macrophages, we utilized scRNA-seq and single nuclei-sequencing (snRNA-seq) from 4 Covid19 cases, 3 LRTD cases and 1 non-LRTD case. Integrating 78,000 cells resulted in 16 cell clusters composed of a mixture of immune and stromal cells (Fig.3a).

**Figure 3.**
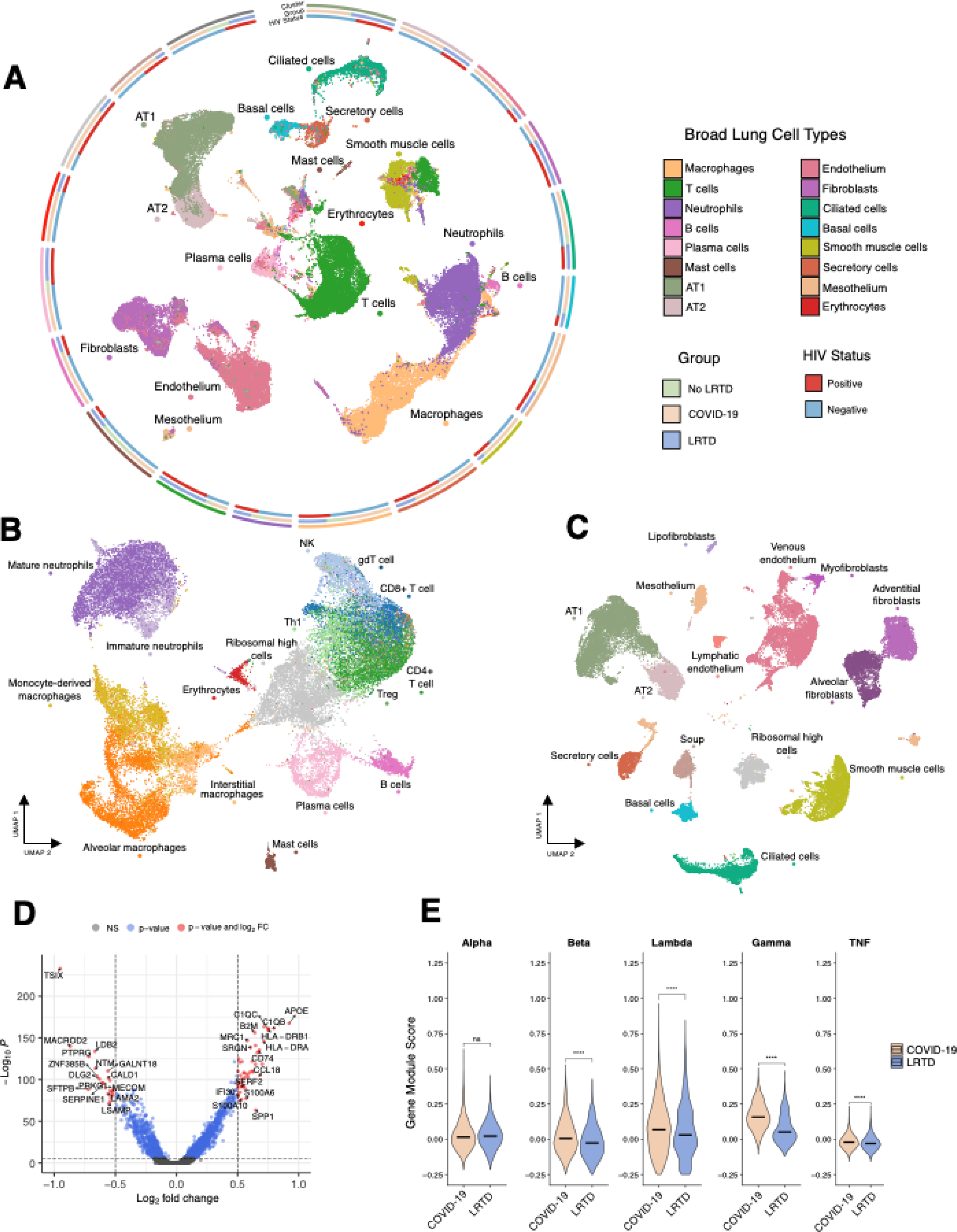
Lung single cell atlas highlights IFN-γ response in alveolar macrophages A) UMAP visualisation of 66,882 lung cells across our cohort, coloured by broad cell types. Outer circos tracks denote the proportion of each cell type across our three disease groups (COVID-19 orange, LRTD blue and No-LRTD green) and whether the cells belong to patients that are HIV positive/negative (blue and red respectively). **B)** UMAP visualisation of 33,504 lung cells reclustered at a higher resolution to characterise the immune landscape, coloured by cell type. **C)** UMAP visualisation of 33,378 lung cells reclustered at a higher resolution to characterise the stromal landscape, coloured by cell type. To note, cells assigned ‘soup’ were not able to be clearly defined by canonical cell type markers and were indicative of multiplets/low quality cells. **D)** Volcano plot showing top differentially expressed genes in alveolar macrophages in COVID-19 compared to LRTD with a significant adjusted p-value (<0.05) and a log-fold change of more than 0.5. **E)** Violin plots showing the gene module score across alveolar macrophages in gene sets associated with the gamma, alpha, beta, lambda and TNF response in COVID-19 compared to LRTD. Black lines indicate the median value across all cells, with asterisks to denote the significance level (ns = non-significant, **** = p <= 0.0001).

SARS-CoV2 transcripts have been detected in scRNA-seq data in other postmortem cohorts^7,8^. We detected few SARS-CoV2 reads suggesting that at time of death there was minimal replicating virus (Extended data Fig.3). This is contrary to our initial prediction of tolerance and viral escape predominating in SSA populations but is consistent with our other data supporting inflammatory rather than direct viral-driven pathogenetic mechanisms.

We then undertook finer annotation of immune (Fig.3b) and stromal/vascular cell pools (Fig3c). We identified alveolar and interstitial macrophages and monocyte-derived-macrophages, consistent with monocyte/macrophage populations identified by IMC (Fig.2a,d). Both mature and immature neutrophils were present. Stromal cells included adventitial and alveolar fibroblasts as well as type I and II pneumocytes (AT1, AT2) and basal, secretory and ciliated epithelial cells. Cell proportions should be interpreted with caution given few cases per group, but they showed cell diversity expansion in the Covid19 and LRTD groups not observed or absent in the LRTD group (Extended data Fig.4a,b).

Principal differences in Covid19 compared to LRTD were in myeloid cells, particularly alveolar macrophages (Fig.3d,e), while few genes were expressed at higher levels in lymphocytes, dendritic cells or stromal cells (Extended data Table 3). In alveolar macrophages top differentially regulated genes included markers of tissue residency (C1QC, C1QB)^31^ and factors shown to mediate lung fibrosis (CCL18)^32^ and apoptosis (S1006)^33^ and activation and recruitment of other myeloid cells (SPP1^34^. IFN-γ response protein (IFI30) and MHC proteins (HLA-DRA, HLA-DRB1) were all upregulated, indicating response to IFN-γ. SPP1 in alveolar macrophages is also linked with smoke-induced lung damage through IFN-γ induction^35^.

This IFN-γ dominant response contrasts with Type I and III dominant interferon responses shown to be critical in pathogenesis in Northern hemisphere Covid19 cohorts^14,36^. Given our IMC data indicating a prominence of alveolar macrophages in the immune response and in alveolar damage we analysed alveolar macrophage interferon response modules: IFN-γ response pathways were strongly upregulated in Covid19 compared to LRTD. IFN-β, IFN-λ and TNF responses were also upregulated but to a lesser degree. Across other myeloid cell IFN responses were heterogenous and TNF response was upregulated in the LRTD group in CD4 T-cells (Extended Data Fig.4c).

### Integration with Human Lung Cell Atlas (HLCA): IFN-γ driven responses in Malawi cohort and type I/III interferon responses in other cohorts

To validate the IFN-γ response in the Malawi cohort compared to type I and III interferon and IL1-dominant responses described in Northern hemisphere cohorts, and to understand the implications for distinct therapeutic approaches we integrated our single cell data with multi-cohort Covid19 (5 cohorts, 60 cases), LRTD (1 cohort, 13 cases) and non-LRTD (23 cohorts, 178 cases) data from the human lung cell atlas^10^ (HLCA)(Fig.4a)(cohorts summarised Fig.1b).

**Figure 4.**
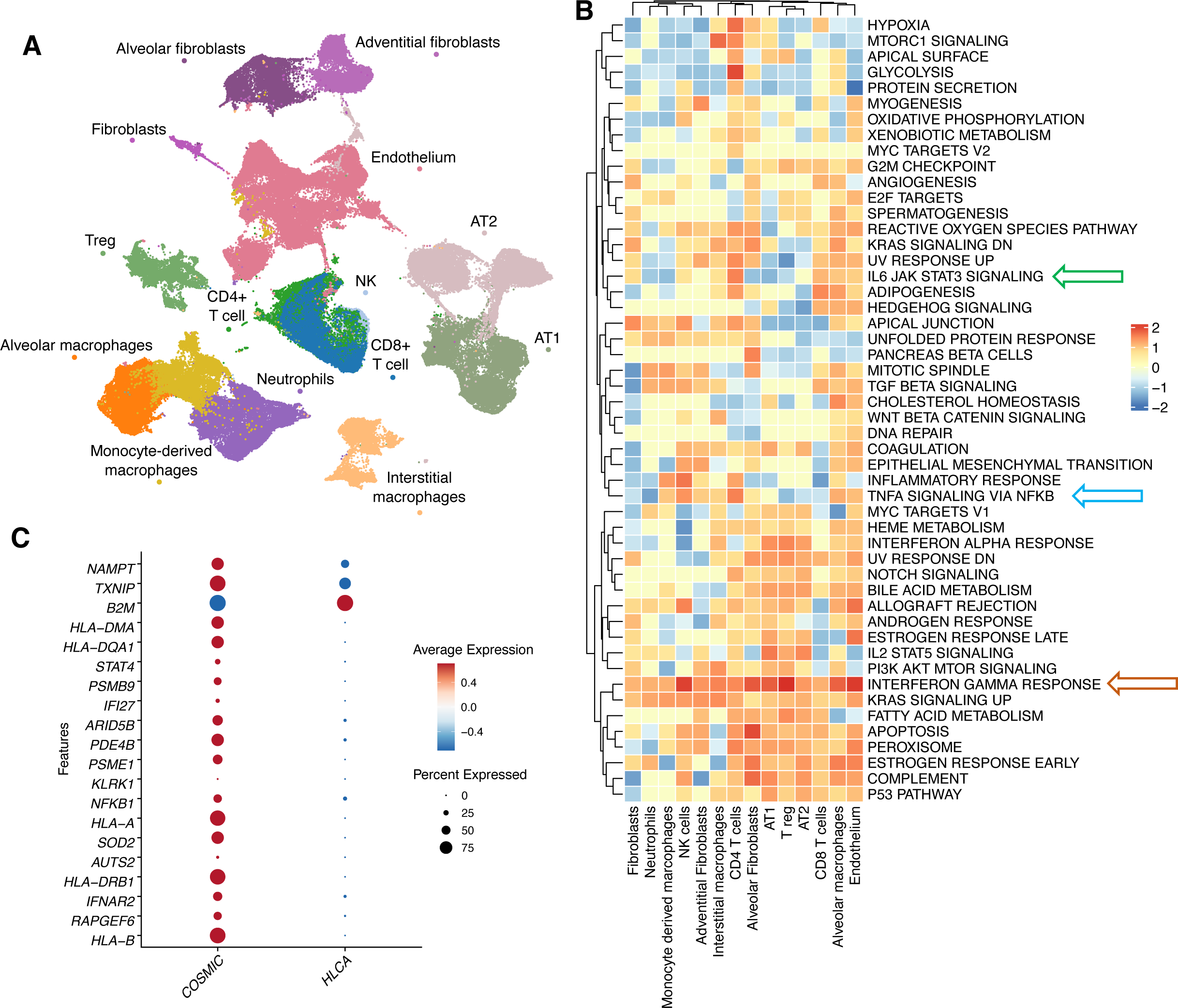
Integration with Human Lung Cell Atlas Covid19 cohorts highlights dominant T-cell macrophage IFN-γ axis in Malawi Covid19 cases A) UMAP visualisation of 147,935 lung cells deriving from integrating cells from Covid19, LRTD and non-LRTD cases from our cohort with cells from the HLCA from non-LRTD, LRTD and Covid19 cases. Clusters are coloured by cell type. **B)** Heatmap showing pathway analysis for differentially expressed genes in our COVID-19 cohort compared to the HLCA COVID-19 cohort. Shown are the 50 canonical hallmark gene sets (for list see Supplemental information) coloured by the normalised enrichment score for each cell type. Gene ontology pathways of interest are indicated by arrows (IL6 JAK STAT3 SIGNALING, green, TNFA SIGNALING VIA NFKB, blue, INTERFERON GAMMA RESPONSE, orange). **C)** Dot plot showing the average expression of top differentially expressed genes in the lung alveolar macrophages that contribute the highest in the hallmark gene set “INTERFERON GAMMA RESPONSE” pathway in our COVID-19 cohort compared to the HLCA COVID-19 cohort.

We used pathway analysis to profile cellular response differences between our cohort and cohorts in the HLCA (Fig.4b). Pathways indicative of IFN-γ response were increased across all cell types in the Malawi cohort (Fig.4b orange arrow). Furthermore, IFNG (IFN-γ gene) was specifically increased in the Malawi cohort in CD4 and CD8 T-cells versus HLCA Covid19 and Non-LRTD groups (Extended data Fig.5), suggesting that macrophages are responding to

**Figure 5.**
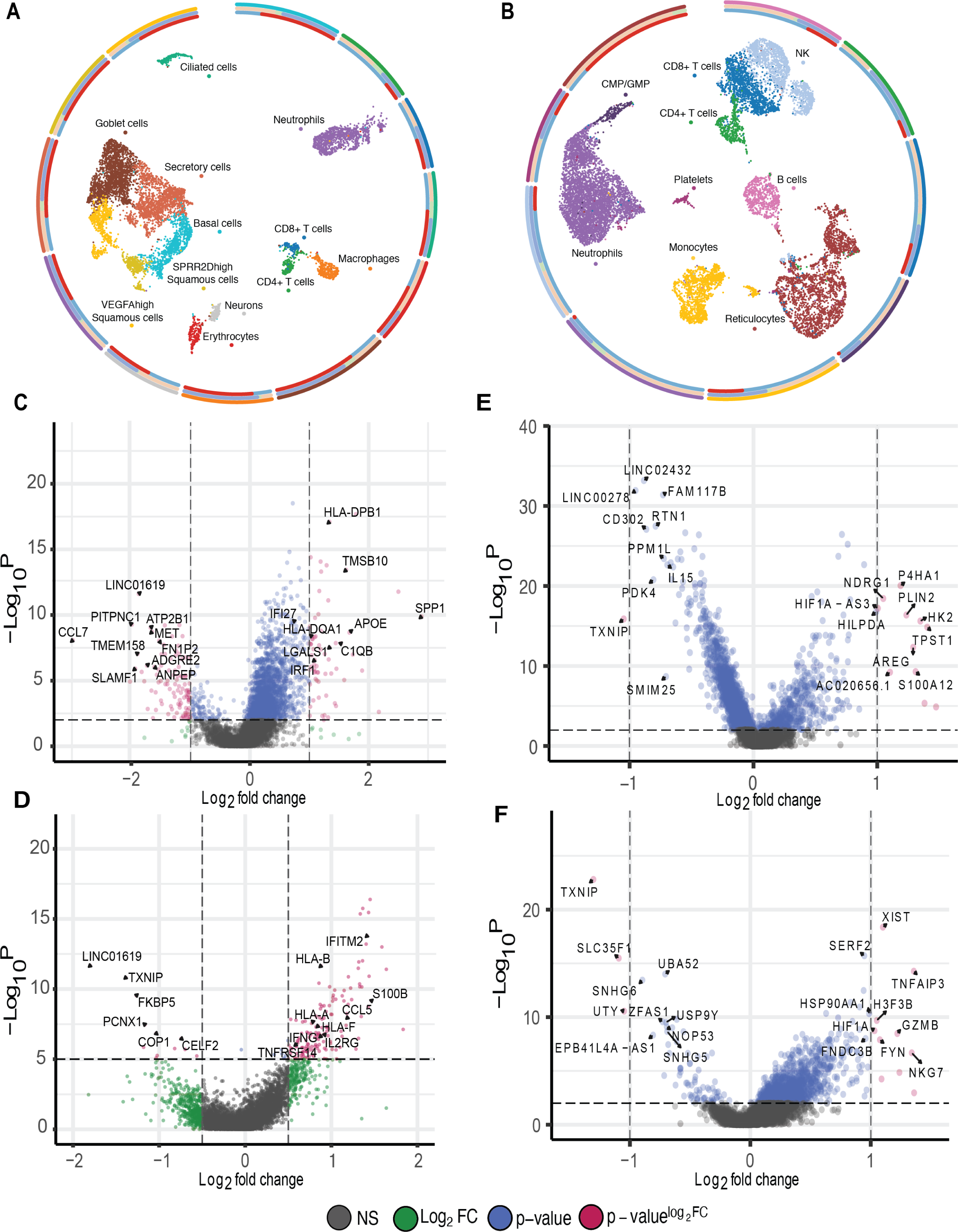
scRNA-seq of nasal and blood cells: nasal but not blood cells parallel lung IFN-γ response. A) UMAP visualisation of 8,098 nasal cells across our cohort, coloured by broad cell types. Outer circos tracks denote the proportion of each cell type across our three disease groups (COVID-19 orange, LRTD blue and No-LRTD green) and whether the cells belong to patients that are HIV positive/negative (blue and red respectively). **B)** UMAP visualisation of 13,350 peripheral blood cells across our cohort, coloured by broad cell types. Outer circos tracks denote the proportion of each cell type across our three disease groups (COVID-19 orange, LRTD blue and No-LRTD green) and whether the cells belong to patients that are HIV positive/negative (blue and red respectively). **C-D)** Volcano plots showing top differentially expressed genes in nasal macrophages and T-cells in COVID-19 compared to LRTD with a significant adjusted p-value (<0.05) and a log-fold change of more than 0.5. **E-F)** Volcano plots showing top differentially expressed genes in peripheral blood monocytes and CD4+ T cells respectively in COVID-19 compared to LRTD with a significant adjusted p-value (<0.05) and a log-fold change of more than 0.5.

IFN-γ produced by T-cells. Other inflammatory pathways showed a mixture of up and downregulation in the Malawi cohort compared to HLCA cohorts, including IL6/JAK/STAT (Fig.4b, green arrow) and TNF-NFKB (Fig.4b, blue arrow), key targets for therapies being used in Covid19. Many of the other interferon-response genes were more upregulated in the HLCA cohorts or had a heterogenous distribution across cells (Extended data Fig.5) although notably monocyte-derived macrophages generally had a higher interferon response in HLCA Covid19 cohorts (Extended data Fig.5).

These data show many shared inflammatory pathways between Malawi and HLCA cohorts but with an amplified IFN-γ response in the Malawi cohort, highlighting IFN-γ production from CD4/CD8 T-cells and response in alveolar macrophages.

### Single-cell analysis of nasal cells may be a useful proxy for lung parenchymal responses

While lung is the principal organ involved in severe and fatal Covid19 disease, lung samples are not easily accessible during life. For future Covid19 waves or other emerging diseases it would be invaluable to predict lung responses using nasal or blood samples that can readily be obtained.

We performed scRNA-seq on nasal cells in 8 cases (5 Covid; 2 LRTD and 1 non-LRTD) and peripheral blood mononuclear cells in 7 individuals (4 Covid19, 2 LRTD and 1 non-LRTD). We recovered 8,098 nasal cells which mapped to ten clusters composing immune and stromal cells and 13,350 blood cells (Fig.5a,b). Nasal macrophages had several similar differentially expressed genes in the Covid19 versus LRTD cases that mirrored lung alveolar macrophage responses including SPP1 and C1QB, genes indicative of proliferation (LGALS1, TMSB10) and MHCII genes (HLA-DPB1, HLA-DQA1) (Fig.5c). HLA-D upregulation is a canonical response to IFN-γ^37^ and consistent with this there was IFNG (IFN-γ gene) upregulation in T-cells in the Covid19 cases in comparison to LRTD cases (Fig.5d). Pathway analysis showed higher levels of IFN-γ response in macrophages and T-cells, further validating an IFN-γ response in these cells (Fig.5e). Blood monocytes in Covid19 versus LRTD cases had upregulation of the alarmin S100A12 and of genes involved in inflammation (AREG) and vascular damage (NDRG1) but not in genes indicative of IFN-γ response and IFNG was not upregulated in T-cells (Fig.5f). Hence in our small cohort nasal cells better paralleled lung response than blood cells, supporting previous Covid19^38,39^ and non-Covid19^40^ studies that highlighted the utility of nasal cells for understanding respiratory immune responses.

Since scRNA-seq is not available in most setting we assessed the extent to which cytokine responses (Luminex) in plasma or nasal fluid could distinguish the inflammatory or IFN-γ response in Covid19 versus LRTD cases. In nasal fluid there was a trend towards several cytokines being higher in Covid19 cases than in LRTD cases, but none significant and no clear difference for IFN-γ (Extended data Fig.6a). There was no clear blood circulating cytokine response pattern, and no circulating cytokine levels were significantly higher in Covid19 compared to other groups (Extended data Fig.6b). A pseudobulk sequencing approach in blood, nasal and lung cells also did not distinguish a clear IFN-γ or any other specific inflammatory cytokine signature between Covid19 and LRTD cases (Extended data Fig.6c-e). Single-cell methods identified an interferon signature and T-cell-macrophage axis, bulk cytokine and gene expression approaches did not. Given very small numbers per group this is perhaps unsurprising. It may stem from greater discriminatory power of single-cell methods and is supportive of the value of single-cell approaches, particularly in small cohorts.

### Stromal cellular interactions are driven by macrophages and vascular interactions by neutrophils

To validate our findings of the role of IFN-γ responding alveolar macrophages in lung parenchymal pathology and neutrophil interactions in vascular pathology, and to predict novel molecular interactions to target therapeutically we used cell interaction methods. First, unbiased receptor-ligand analysis of our scRNA-seq data highlighted that a large proportion of the imputed interactions in the lung involved alveolar macrophages, interacting with fibroblasts, epithelial cells and other immune cells including CD4 T-cells (Fig.6a), in keeping with our findings from cell proportions, cellular histology and scRNA-seq.

**Figure 6.**
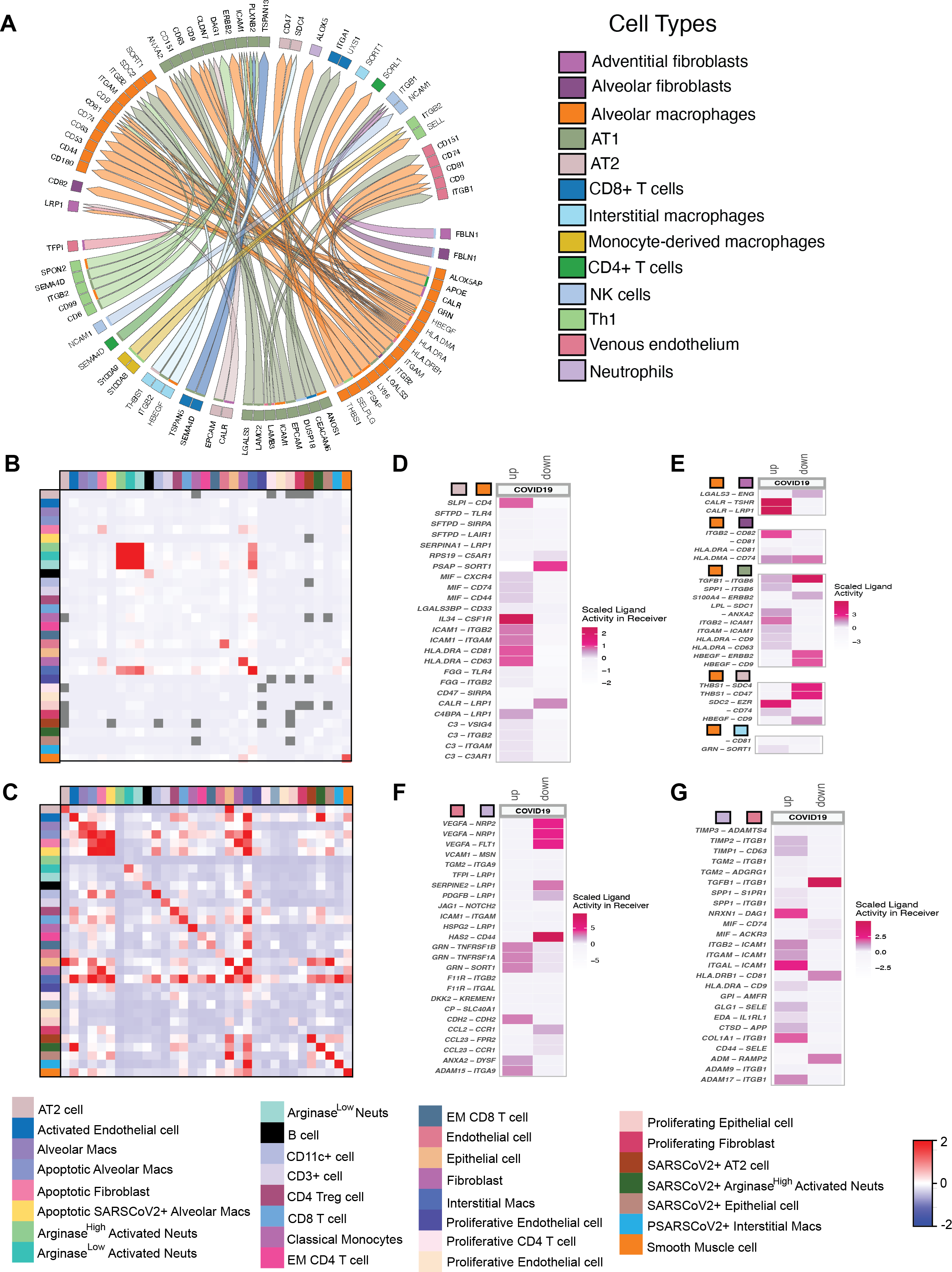
Spatially resolved cell interaction analysis predicts molecular mechanisms of alveolar and endothelial pathology A) Circos plot showing the top 50 differentially expressed interactions upregulated in our COVID-19 cohort compared to LRTD. Segments are coloured by cell type with ligands and receptors labelled on the outside. Direction of the arrows show the senders of communications i.e. expression of ligand, and receiver of communications. Inner tracks on sender segments are coloured by the receiving cell type for ease of interpretation. **B-C)** Heatmaps showing co-localised cell types as shown by the IMC providing insight into potentially interacting cell types in the lung. **D)** Heatmap showing up/down-regulated interactions in COVID-19 compared to LRTD driven by AT2 pneumonocytes to alveolar macrophages. Coloured boxes indicate cell type with the ligand-expressing cell type followed by the receptor-expressing cell type. **E)** Heatmap showing up/down-regulated interactions in COVID-19 compared to LRTD driven by lung alveolar macrophages to lung epithelial cells and interstitial macrophages. Coloured boxes indicate cell type with the ligand-expressing cell type followed by the receptor-expressing cell type. **F)** Heatmap showing up/down-regulated interactions in COVID-19 compared to LRTD driven by lung endothelium to neutrophils. Coloured boxes indicate cell type with the ligand-expressing cell type followed by the receptor-expressing cell type. **G)** Heatmap showing up/down-regulated interactions in COVID-19 compared to LRTD driven by neutrophils to lung endothelium. Coloured boxes indicate cell type with the ligand-expressing cell type followed by the receptor-expressing cell type.

To validate these interactions in a spatial context we used neighbourhood enrichment analysis of IMC data to identify cells located close to each other with greater than expected frequency as an indicator of their likelihood to be interacting (Fig.6b,c). In the non-LRTD there were no significant interactions (Extended Data Fig.7). The LRTD group was completely dominated by neutrophil interactions (Fig.6b). In the Covid19 group several neighbourhood enrichments were prominent – principally alveolar macrophages (with and without SARS-CoV2-S and apoptosis) with apoptotic fibroblasts and to a lesser extent type II pneumocytes (Fig.6c). This supports the role in pathogenesis of alveolar macrophages including the apoptotic population present in Malawi but not USA or Brazil cohorts. In contrast the most prominent neighbourhood enrichment for neutrophils was between SARS-CoV2-S+,Arg^high^ neutrophils and activated endothelial cells implicating neutrophils in endothelial/vascular pathology.

We then looked at validated interactions in Covid19 in closer detail in scRNA-seq data. Macrophage interactions were frequently from ligands on type II pneumocytes to receptors on alveolar macrophages (Fig.6d), in keeping with type II pneumocytes cells generally being a principle infected cell^41^. Several of these interactions involved macrophage inhibitory factor (MIF) from type II pneumocytes with CD74, CD44 and CXCR4 on macrophages, a classical response chain in macrophages and a key initiator of proliferation, chemotaxis and activation^42^. ICAM-1 on type II pneumocytes was predicted to signal to integrins (ITGB2-ITGAM) on alveolar macrophages, an interaction involved in cellular attachment during recruitment. Another strong predicted interaction was IL-34-CSF1R, involved in triggering macrophage activation and chemotaxis. Reciprocally there were several interactions between alveolar macrophages and epithelial cells consistent with our IMC data that indicate their role in alveolar pathology. These included SPP1 and TGFβ with type II pneumonocyte integrins (ITGB6) (Fig.6e), interactions implicated in lung pathology and fibrosis^34,35,43^. We identified multiple neutrophil interactions with endothelial cells indicating processes involved in neutrophils attachment to the vascular wall (e.g., ITGAL-ICAM-1) and of activation by neutrophil granule proteins (GRN-TNFRSF1A) (Fig.6f,g), providing molecular validation supporting their role in coagulation, endothelial activation and vascular pathology indicated by IMC.

These data highlight the value of a combined scRNA-seq and IMC approach. They provide spatial and receptor-ligand validation for roles of alveolar macrophages in molecular processes that are plausibly involved in alveolar damage and lung fibrosis, and for neutrophils in endothelial activation. The data predict specific molecular interactions involved in these processes. If validated by further work, some of these interactions may be plausible targets for intervention, e.g., MIF for which several small molecules are in clinical development for therapy in inflammatory disorders^42^.

## Discussion

We conducted minimally invasive autopsies on fatal Covid19 and other LRTD and non-LRTD cases in a Malawian population and characterised pulmonary, blood and nasal immune responses using scRNA-seq and IMC. While other studies have used these techniques for Covid19 investigations in other settings, this is the first such study in a SSA population and to our knowledge some of the first scRNA-seq data from lung samples in any SSA population. Our de-identified data are provided open access, including tools for visualising single-cell and histology data, making an important resource for furthering the global understanding of Covid19 pathogenesis, immune responses in SSA populations and more widely for the human cell atlas.

Given that many parasitic infections induce immune tolerance we hypothesised that there might be an attenuated immune response in SSA populations, blunting immune-mediated viral clearance and leading to high viral-loads in individuals who present with life-threatening disease. If so, pathology might be driven by direct-viral effects rather than hyperinflammation, indicating a need for different treatment approaches from Northern hemisphere cohorts. In fact, we found a robust immune response and comparatively low levels of virus, surprisingly even in highly immunosuppressed cases with HIV. Our data indicate that pathology is driven by inflammation, with many similarities to other non-African cohorts, in both histopathology and immunology. These similarities are reassuring, indicating that many principles for diagnosis and treatment can likely be extrapolated from more extensively investigated populations. However, there were also differences that may have implications for therapy, in particular IFN-γ responses were upregulated in comparison to a large multi-country integrated HLCA dataset. IFN-γ was produced by T-cells, with alveolar macrophages the principle responding cells. Using IMC, we showed that these resident macrophages were the dominant cell in pathological lesions. Spatially-resolved IMC interaction analysis and scRNA-seq receptor-ligand analysis orthogonally validated these processes. In contrast IL6 and TNF responses were not as prominent. scRNA-seq of nasal cells also identified IFNG upregulation in T-cells and evidence of IFN-γ response in macrophages in a sample type that is readily accessible, supporting prior data on the utility of nasal cells as an accessible proxy for lung responses.

There is cross-over between the responses of different interferons and IFN-γ signal has been detected previously in Covid19 lung^7^, yet it is interesting to consider why there was such a marked upregulation in our cohort compared to the large integrated HCLA dataset. IFN-γ response has been shown to be a key component of effective immunity to malaria and is augmented in malaria exposed individuals, in part through epigenetic changes termed trained immunity^17^. Increased IFN-γ response was a key difference in SSA (Gabon) versus European individuals exposed to controlled human malaria infection and a correlate of protection^16^. While type I/III interferons are more typically involved in clearance of SARS-CoV2 and other respiratory viruses^44^, IFN-γ also plays a role, particularly in macrophages^45^. Considering our data with these prior studies we propose that trained immune responses to prior infections may favour an accelerated macrophage IFN-γ response. We hypothesise that this may be a double-edged sword in Covid19 in SSA: such an accelerated trained response may generally be protective (through more rapid viral clearance), but in a subset of patients it may lead to accelerated hyperinflammation and collateral tissue damage. This hypothesis is supported by the short time between symptom onset and death in our cohort already with a clear hyperinflammatory response. Further exploration of macrophage responses in both SSA and non-SSA populations is therefore warranted.

Considering the potential for translation the existing therapies for Covid19 target JAK/STAT (Barcitimib), IL6 (Toculimazab/sarilumab) or TNF (infliximab)^14,15^. JAK/STAT signalling is a conserved pathway for interferon responses including IFN-γ^45^. Thus our data, if corroborated, support potential efficacy of Baricitinib over other treatments. Barcitinib is a small molecule (tablet) and thus highly suited to wide distribution^15^.

Our data have several limitations. Our cohort was small and in a single centre. Although single-cell methods have a higher capacity to resolve complex data in small sample sizes, many analyses in our study were underpowered. It is thus unclear how representative our data are of the wider Malawi or other SSA populations. Studies in other settings and ideally large multi-centre studies, are needed. While this would be a complex undertaking, we have demonstrated that single-cell methods are feasible in a SSA setting, and our study provides useful templates. While lung samples cannot readily be obtained in live patients, postmortem studies have limitations: cells may change or degrade; pathological processes present early in disease are likely missed. Yet, postmortem studies in northern hemisphere settings with longer postmortem intervals identified validated targets^7^. While minimally invasive autopsy is more feasible and acceptable than traditional open autopsy, blind sampling may attenuate the identification and sampling of areas of pathology. However, except for large airway pathology, which was not sampled, most Covid19 features were identified. The studies that we used for comparisons had significant variation in methods and demographics from ours which may induce noise and bias. We used data-integration methods which reduce but do not eliminate this. Reassuringly findings were validated both by comparison to Human Lung Cell Atlas and orthogonal IMC data.

Our data establish the feasibility and utility of single-cell analyses in postmortem studies in a SSA setting and provide a resource of lung, blood and nasal cells alongside histology and spatial characterisation on a population not yet represented in the human cell atlas. Our work indicates differences in the Covid19 inflammatory response among a Malawian cohort compared to other non-African cohorts, highlighting IFN-γ as a potential target for intervention and supporting the utility of scRNA-seq of nasal samples to assess respiratory-tract immunology in LRTD.

## Online Methods

### Ethics and sensitisation

The study protocol was approved by the National Health Scientific Research Committee (NHSRC) in Malawi (Protocol number 07/09/1913) and by the Medical Veterinary Life Sciences ethics committee in Glasgow (protocol number 200190041). Informed consent was taken from the families of deceased patients. We underwent a full sensitisation process for the study with all staff on the recruiting wards in our hospital to discuss the study and consider the best way of sensitively conducting recruitment and informed consent. This work was led by two social scientists (L.S., D.N.) one with specialised in Bioethics (D.N.). Details of our approach and considerations for recruitment are published separately (*In press*).

### Patients

We recruited patients aged 45-75 admitted to Queen Elizabeth Central Hospital, Blantyre between October 2020 and July 2021 during which there were two epidemiological waves driven by different SARS-CoV2 variants: Beta (Dec 2020-Feb 2021) and Delta (May-July 2021). Patients admitted with respiratory signs were routinely tested for SARS-CoV2 at QECH. We recruited cases into three groups based on clinical criteria: 1) a Covid19 group (n=9) with clinical features suggesting acute respiratory distress (ARDS, oxygen requirement and either respiratory signs on clinical examination or chest x-ray changes or both) and who had at least one nasal swab positive for SARS-CoV2 on admission; 2) A non-Covid19 LRTD (lower respiratory tract disease) group (n=5) who had clinical signs of ARDS but were negative for SARS-CoV-2 on admission and during hospitalization; 3) a no LRTD, COVID-19 negative group (n=2) who had no oxygen requirement and no clinical signs of LRTD and for whom the admission and any subsequent nasal swabs were negative for SARS-CoV2 on PCR (Fig.1b, Extended Data Table 1). The study only recruited cases who died between 12 midnight and 12 noon to minimise the postmortem interval and to avoid doing any autopsies at night.

### Minimally invasive autopsy

We used minimally invasive sampling to conduct autopsies with large-bore needle biopsies of organ samples rather than full autopsy^19^. Being more culturally acceptable, MITS is widely used to determine cause of death in paediatric studies^19,46-48^, showing good concordance with full autopsy^46^. From our ongoing paediatric MITs studies in Malawi, we adapted protocols for adult patients with Covid19 to obtain tissue suitable for single cell RNA-sequencing (scRNA-sequencing) and IMC. Based on CHAMPS but with adaptations. In particular, a larger calibre needle (11 gauge) was used for biopsies to obtain larger tissue samples. Samples were taken from the brain from supraorbital sampling from both left and right sides. From each lung samples were taken from lower middle and upper zones from a single entry-point, angling the needle to sample different areas. Nasal cells were collected from the nasal inferior turbinate using curettes (ASL Rhino-Pro, Arlington Scientific). Two curettes were collected from each nostril and the cells placed immediately into ice cold Hypothermosol (StemCell). Cells were transported on ice in a cold box immediately to the lab and were spun at 300g for 5mins either for immediate processing for scRNA-seq or were stored in Cryostor 10 (see below). Nasal fluid was collected using matrix strips (Nasosorption, Hunt Developments). One strip was used per nostril.

PPE was worn by all staff involved in the autopsies and for all work in the lab. Lab work on samples was done in vented laminar flow hoods.

### Processing of Storage of samples

Biopsies from each organ were collected in three different ways for different downstream workflows: 1) for paraffin embedding for histology and IMC put in 10% neutral buffered formalin, 2) for viable cells put in ice cold hypothermosol (StemCell) for transport to the lab and then slow frozen in Cryostor 10 (StemCell), 3) for snap frozen cells put in cryovials which were then sealed and immediately submerged in liquid nitrogen.

Biopsies were fixed in 10% neutrophil buffered formalin for 4 - 8 hours, rinsed in water and then embedded in paraffin blocks. Samples for viable cells were rinsed and cut into pieces of approximately 20mm – 50mm and then put into ice cold cryostor for 15 – 30 mins before transfer to a -80°c freezer in a chilled cryogenic storage container (CoolCell, Corning).

Blood cells collected into sodium heparin tubes were separated from plasma by spinning at 400g for 10 minutes. Plasma was then removed and spun for an additional 10 minutes at 1500g and plasma frozen in aliquots at -80°c. Cells were resuspended in 10% fetal bovine serum in PBS and PBMCs were separated using ficoll histopaque with a 27min spin at 450g and either used immediately for scRNA-seq or pelleted and resuspended in ice cold Cryostor 10 and then moved to a -80°c freezer in a chilled cryogenic storage container (CoolCell, Corning). The following day samples were moved from the -80°c freezer to liquid nitrogen for long term storage. Snap frozen samples were transferred in a liquid nitrogen dewar and then moved to liquid nitrogen storage tanks for long-term storage.

### Pathology and organ specific scoring

Formalin-fixed tissues were paraffin embedded (FFPE) for lung, bone marrow, brain, spleen, and liver to make blocks. FFPE blocks were sectioned at 2-4 μm thickness, mounted on glass slides and stained with haematoxylin and eosin (H&E). A medical pathologist (S.K.) reviewed tissue slides, alongside patients’ history and antemortem lab results, per standard clinical practice and also completed an organ specific scoring proforma that included Covid19 features (Supplemental table 1). Then, for a non-biased assessment, two additional pathologists, blinded to diagnosis, scored the lung pathology in all patients using systematic scoring criteria^49^. Lung tissue was scored independently by two additional pathologists (C.A. and V.H.), who were blinded to patient history and previous diagnoses. Following individual scoring, any discrepancies were agreed by joint review of the slides until a consensus was reached. The lung scoring was semiquantitative for the parameters indicated in Extended data, Fig.1a-c. Subsequently, we characterised each sample with a dominant histological characteristic, e.g., fibrinopurulent inflammation/pneumonia in case the neutrophil infiltration with fibrin-extravasation was marked next to a mild infiltrate of lymphocytes, plasma cells and macrophages. Whole tissue slides from lung samples in our 9 Covid19 cases can be accessed in their entirety, and visualised at various magnifications, as if they were observed under a microscope using our virtual microscope tool: https://covid-atlas.cvr.gla.ac.uk (de-identified slides will be uploaded and publicly viewable on publication).

After scoring, in each lung biopsy, the most representative areas were manually selected based on the scoring performed on the H&E-stained section to create the tissue microarrays (TMA) with cores of 1mm in diameter using the TMA Grand Master (3Dhistech, USA) and CaseViewer software (version 2.4.0119028). At least 8 regions of interest were taken from each case (4 left, 4 right). From the newly created TMA-FFPE-blocks, 4 μm thick sections were cut and used for downstream imaging mass cytometry (IMC).

### Cause of death attribution

A panel consisting of the pathologist who reviewed the cases, respiratory physician, intensive care physician, infectious disease physician and two trainee doctors reviewed all the cases to assign a cause of death. Codes assigning death were given according to ICD codes and using the standard coding system used for death certification. The review consisted of a review of the clinical notes, pre and post mortem lab results and the pathology report. Each member reviewed the documents independently and reached an individual verdict. When there was discrepancies a consensus was reached through discussion.

### Multiparameter Cytokine Assay

Cytokine levels were measured in plasma and nasal fluid samples using Luminex with the Inflammation 20-Plex Human ProcartaPlex™ Panel (Thermofisher, EPX200-12185-901) according to the manufacturers protocol and levels measured with a Luminex MagPix device. Data were transformed with a log2 and for the visualistion with ComplexHeatmap in R with a Z-score by gene. For the statistical tests of genes associated with the IFN-y pathway we used a Welch Two Sample t-test. No significant differences for those genes was found between the Covid-19 and LTRD samples.

### Imaging mass cytometry (IMC)

Sections from tissue micro arrays underwent deparaffinisation, followed by antigen retrieval at 96°C for 30 min in Tris-EDTA at pH 8.5. Non-specific binding was blocked with 3% bovine serum albumin for 45 min, followed by incubation with lanthanide-conjugated primary antibodies (overnight at 4°C) which were diluted in PBS with 0.5% BSA (Table of antibodies in Supplemental information). Antibodies were conjugated with metals using Maxpar Antibody Labeling Kits (Standard BioTools) and were validated with positive control tissue (tonsil and spleen for immune-targeted antibodies). Slides were then washed with 0.1% Triton-X100 in PBS, followed by nuclear staining with iridium (1:400, Intercalator-Ir, Standard Bio Tools) for 30 minutes at RT, and finally briefly (10 s) washed with ultrapure water and air-dried. Images were acquired on a Hyperion imaging mass cytometer as per manufacturer’s instructions (Standard BioTools). Each TMA core was imaged in a separate region of interest.

### Imaging Mass Cytometry (IMC) analysis

Preprocessing, imaging denoise, cell segmentation and extraction of single-cell features were performed using a combination of Python and R packages, such as ImcSegmentationPipeline, IMC-Denoise^50^ and DeepCell^30,51^ as described previously^13^. Briefly, for the single-cell analysis, the annotated data object was generated, protein expression raw measurements were normalized at the 99th percentile to remove outliers. In Scanpy (Single-Cell Analysis in Python v 1.9.1)^52^ PCA, batch-correction and harmony data integration was performed to compute and plot the UMAP embeddings (umap-learn Python package, v 0.5.3). Next, automated cell type assignment using the Python package Astir (ASsignmenT of sIngle-cell pRoteomics v 0.1.4)^51^, was applied to identify the major cell types expected to be found in the lung tissue according to the antibody panel utilized. For cell assignment with Astir, the following information to label cells based on a broad ontogeny (metaclusters and major cell types) and the proteins (lineage markers) to be most expressed in each expected cell type were used. Metaclusters and major cell types: (a) myeloid: macrophage, neutrophil; (b) lymphoid: CD8 T cells, CD4 T cells, B cells; (c) vascular: Endothelium, red blood cells (RBCs); (d) stromal: fibroblast, smooth muscle cell, epithelial. Cell types: (a) macrophage: CD163, CD206, CD14, CD16, CD68, CD11c, Iba1; (b) neutrophil: CD66b, Arginase1; (c) CD8 T cells: CD3, CD8; (d) CD4 T cells: CD3, CD4; (E) B cells: CD20; (f) endothelium: CD31; (g) fibroblast: Collagen1; (h) smooth muscle cell (SMC): SMA; epithelial: PanCK; RBCs: CD235ab.

After cell assignment, cells labelled as “other” or “unknown” were filtered out from downstream analysis, the annotated data object was subset into the major cell types identified, i.e., macrophages, neutrophils, lymphoid, vascular, epithelial and stromal and Phenograph Louvain clustering (with 200 nearest neighbors)^53^ was performed for each cell population separately using a small set of specific lineage marker and functional proteins, as previously described^13^. The finer cell type annotation was used to evaluate the frequency and absolute counts of cell types across clinical groups, histopathological lesions and HIV status. Differential abundance analysis was also performed using the scanpro and scCODA Python’s packages^54^ and miloR R package (v 1.4.0)^55^. Spatial statistics analysis based on the coordinates of the cells in the ROIs, were performed using the Python packages Squidpy (Spatial Quantification of Molecular Data in Python v. 1.2.2)^56^. Building of the spatial graph, calculatin and plotting of the neighborhood enrichment scores were performed as previously described^13^.

### Integration of Malawi IMC data with other available IMC COVID-19 lung data

IMC COVID-19 data from post-mortem lung samples from a Brazilian^13^ and US^30^ fatal cohort were integrated with the Malawi IMC dataset. First, datasets were concatenated in Scanpy taking the “inner” (intersection) of all common protein markers in the panels across the 3 IMC datasets. Then, with scvi-tools^57^ we applied different integration methods, such as harmony and variational autoencoder (VAE)-based methods, such as scVI and scANVI. Analysis of the UMAP embedding of the integrated versus non-integrated data showed that harmony and scANVI performed better and in downstream analysis we used harmony-integrated output. Next, cell identities were standardized (label harmonization), which refers to a process of checking that labels are consistent across the datasets that are being integrated. Finally, cell frequencies in the post-mortem lung across all 3 cohorts were plotted and differential abundance analysis was performed using scanpro (https://github.com/loosolab/scanpro) and scCODA Python’s packages^54^ and miloR R package (v 1.4.0)^55^.

### Dissociation of lung cells from frozen samples and single nuclei preparation

Lung samples were dissociated both from fresh samples and from slow frozen samples that had been stored in liquid nitrogen. Slow frozen cells were defrosted using a defrosting protocol described previously^7^. Fresh or defrosted frozen cells were then dissociated adapting methods developed previously^58^. Briefly cells were dissociated in a buffer containing 400μg/mL of Liberase DL (Sigma), 32U/ml of DNAse I (Roche) and 1.5% BSA in PBS (without calcium and magnesium). The tissue was put in buffer (4 times weight:volume) in a GentleMACS C-tube *(Miltenyi 130-096-334)* minced with scissors and then run on a GentleMACS dissociator (*130-093-235) on* programme “C-lung 01_02”. Dissociation was achieved by warming tissue on an orbital shaker in a chamber at 37C for 30 mins and running “C-lung 01_02” twice more; once at 15 mins and once at 30mins. Enzyme was neutralised by diluting with 10ml of ice cold 20% FBS with 32u/ml of DNase and the sample was filtered through a 100μM strainer (352360) and samples were subsequently kept on ice with all centrifuge and antibody incubation steps at 4c. Cells were pelleted by spinning at 300g for 5 mins and red cells removed by incubation with ACK buffer for 5mins. For frozen cells debris were removed using a debris removal solution (Miltenyi, 130-109-398) according to the manufacturers protocol. Single nuclei were prepared from snap frozen lung samples as described previously^7^. Briefly, frozen lung tissue was kept on dry ice/ liquid nitrogen until processing was started. Tissue was added to a gentle MACS C-tube containing 2ml of freshly prepared nuclei extraction buffer which contained RNAse inhibitors; 0.2 U/µL RNaseIN Plus RNAse inhibitor (Promega) and 0.1 U/µL SUPERasin RNAse inhibitor (Thermofisher scientific). Dissociation was achieved by running the C-tube on GentleMACs dissociator on program ‘’m_spleen_01’’ for 1 minute. The sample was then filtered using a 40 µM strainer. The C-tube and strainer were rinsed using a buffer containing 0.1% enzymatics RNAse inhibitor (Enzymatics). Sample was then pelleted by spinning at 500g for 10 minutes at 4^0^C. Pellet was then resuspended in 500ul of 1xST without RNAse Inhibitor. The sample was then filtered using 35µM strainer, a 10 µL volume was loaded on haemocytometer for counting.

### Single cell and single nuclei partitioning and library preparation

10x 3’ 3v chemistry was used for all samples. For fresh lung samples we loaded 10,000 cells into one channel of a 10x chip (1000120). For fresh nasal and blood samples we labelled the nasal and blood samples with different hashtags and pooled them at a 1:1 ratio and loaded 10,000 – 20,000 cells. For frozen nuclei and single cell samples we pooled samples from 3 – 6 different cases aiming for equal ratios and loaded 20,000 – 40,000 cells/nuclei. Libraries were prepared according to the manufacturers protocol and sequenced with an Illumina NextSeq2000. To make these data available for analysis by others, reads were submitted to ArrayExpress (E-MTAB-13544).

### Analysis of single cell data

The reads were mapped using Cellranger (version 7, including introns) to a combined reference of human, Covid19. For mixed genotypes, samples were separated using SoupOrCell^59^, and for hashtag oligonucleotide labelled mixes of nasal and blood samples from the same patient, the CiteSeq protocol was followed. Data analysis was conducted in R version 4.2.1, utilizing Seurat^60^ for data integration within each tissue and with the Human Lung Cell Atlas (HLCA) using Harmony. Differential expression analysis was performed with MAST, and cell-cell interactions were assessed using multinichenetR (https://www.biorxiv.org/content/10.1101/2023.06.13.544751v1). All figures were generated in R. The different datasets are accessible on the Glasgow Cell Atlas website: http://cellatlas.mvls.gla.ac.uk/. See additional details in supplemental information.

### Funding and Acknowledgements

We would like to thank families who agreed to take part in this research and all health care professionals at Queen Elizabeth University Hospital who cared for the patients during life, supported bereaved families and helped with recruiting patients and morticians who helped with autopsies. We thank Brigitte Dennis and staff at the Core laboratory at Malawi Liverpool Wellcome for help with processing laboratory samples, Dr Gareth Howell and the University of Manchester Flow Cytometry Core Facility for helping with the IMC investigations and David McGuinness and Julie Gilbraith and other staff at Glasgow Polyomics for technical advice and help with sequencing of single-cell and single nuclei libraries, Monica Soko and Karl Seydel at Blantyre Malaria Project laboratory at KUHES for running the Luminex samples. We would like to thank Alex Shalek, Sergio Heli Triana Sierra and Chloe Villani for technical advice and Alex Shalek for shipping reagents to Malawi when we could not obtain them from our suppliers. We thank Megan McLeod for suggestions on the manuscript. This work was funded through a grant to C.A.M. from the Bill and Melinda Gates Foundation (INV-018138). C.A.M. is a UKRI Medical Research Council (MRC) MRC Future Leaders Fellow (MR/V025856/1). K.N.C. was supported by the MRC (MR/R010099/1). MR/R010099/1 was jointly funded by MRC and the UK Department for International Development (DFID) under the MRC/DFID Concordat agreement and was also part of the EDCTP2 program supported by the European Union. M.M. is supported by a Wellcome Center award (number 104111). J.LS.F. was supported by the Sao Paulo Research Foundation (FAPESP grant 2019/015782 and 2016/12855-9) and it is currently supported by MRC (MR/W018802/1). F.T.M.C is supported by the Sao Paulo Research Foundation (FAPESP grants 2020/05369-6 and 2017/18611-7). W.M.M. is supported by the Amazonas Research Foundation (FAPEAM N. 005/2020 - PCTIEMERGESAÚDE – AM). M.V.G.L., G.C.M., W.M.M. and F.T.M.C. are CNPq research fellows. The University of Glasgow Wellcome Centre for Integrative pathology (104111) and Malawi-Liverpool-Wellcome Programme are supported by Core funding from Wellcome. This publication is part of the Human Cell Atlas –– www.humancellatlas.org/publications/

## Data Availability Statement

scRNA-Seq: Raw data and processed count matrices are deposited at the EBI ArrayExpress (Accession number E-MTAB-13544 (private until publication). Fully processed. RDS objects of the scRNA-seq analysis and IMC can be found through the GitHub repository (see Code Availabilty Statement). The atlases are browsable using the cellxgeneVIP platform hosted by the University of Glasgow at the following URLs:

Lung Atlas - https://cellatlas-cxg.mvls.gla.ac.uk/COSMIC/view/COSMIC_Lung_Atlas.h5ad/

Lung Immune Atlas - https://cellatlas-cxg.mvls.gla.ac.uk/COSMIC/view/COSMIC_Lung_Immune_Atlas.h5ad/

Lung Stromal Atlas - https://cellatlas-cxg.mvls.gla.ac.uk/COSMIC/view/COSMIC_Lung_Stromal_Atlas.h5ad/

Nasal Atlas - https://cellatlas-cxg.mvls.gla.ac.uk/COSMIC/view/COSMIC_Nasal_Atlas.h5ad/

Blood Atlas - https://cellatlas-cxg.mvls.gla.ac.uk/COSMIC/view/COSMIC_Blood_Atlas.h5ad/

Histopathology slides on virtual microscope: https://covid-atlas.cvr.gla.ac.uk

Metadata for the cases (without identifying information) is provided in Extended Table 1. IMC - https://cellatlas-cxg.mvls.gla.ac.uk/COSMIC/view/COSMIC_IMC_Lung.h5ad/

## Code Availability Statement

All R scripts for the scRNA-Seq analysis and figure generation can found at (https://github.com/olympiahardy/COSMIC_Malawi_Covid_Atlas). Python scripts to process the imaging mass cytometry and figure generation can also be found at the above repository.

## Author contributions

Research protocol, Malawi (CAM, VW, DN); Sensitisation, Malawi (LS, TN, MS, DN); Recruitment of patients, Malawi (MS, TN, LS, DN, DC, CE, AT, FZ); Autopsies, Malawi (SK, CA); Pathology report, Malawi (SK); Histopathology scoring and identification of regions of interest (CA, VH); Clinical characterisation and cause of death attribution, Malawi (TN, MS, SK, PB, BM, SL, VW, DD), Lab protocols and processing of tissues, Malawi (JN, CN, FD, WN, TN, SK, CP, CAM), Set up of single cell protocols and optimisation of single cell methods, Malawi (JN, CAM), Single cell experiments (JN, CAM, CN, LN, JP); designed and conducted IMC experiments (MH, KC); Analysis of histology data (CA, VH, CAM) Analysis of single cell data (OH, TO); Protocols, sample collection and processing of Brazil samples (M.F., F.F.AV., M.B., C.C.J.,T.T. and V.S.S); Analysis of IMC data (J.L.); Prepared figures (OH, JL, TO, KB, CAM); Uploaded data onto repositories and interactive modules (OH, TO, VH); Conceptualisation (CAM, TO, VW, DD, SK); Wrote the grant (CAM, TO, DD, VW, BK); Supervision (CAM, TO, MM, DD, VW, KC, SK, MP); Drafted the manuscript (C.A.M.); Edits of the manuscript (JN, OH, JL, KB, KC, MP, DD, MM, TO), all authors reviewed and approved the final manuscript.

## Figure legends –– Extended data

**Extended data Figure 1 |** *The histopathology of fatal Covid19 versus fatal non-Covid19 LRTD and non-LRTD in Malawian cases.*

Histopathology in the left and right lungs of the 16 cases was scored systematically using pre-defined criteria by two pathologists who were blinded to clinical information. We used identical scoring to a Brazil cohort that we have published on separately. A – C are histograms of the scoring: **A**) Comparsion of Covid19 (n=9) and non-Covid19 fatal lower respiratory tract disease (LRTD) cases (n=5). **B**) Comparison of histological features between HIV+ Covid19 cases (n=5) and HIV-Covid19 cases (n=4). **C**) Comparison of Covid19 cases from Malawi cohort (n=9) with cases from Brazil cohort (n=20). A two sided T-test was used to compare lesion frequencies with no correction for multiple comparisions *=p=<0.05. **D**) PCA of cases split by groups. E) UMAP of same data, including HIV status

**Extended data Figure 2 |** *Cell atlas and phenotype of cell types identified in the post-mortem lung tissue determined by Imaging Mass Cytometry (IMC).*

**A**) Phenotype representation of each cell type identified in the lung samples. The heatmap shows the mean expression of each protein marker in the IMC panel in each cell type identified in the post-mortem lung tissue. **B**) Frequency of the immune cell types identified in the post-mortem lung samples by IMC according to clinical groups and according to HIV status within the COVID-19 group. **C**) Frequency of the stromal cell types identified in the post-mortem lung samples by IMC according to clinical groups and according to HIV status within the COVID-19 group. **D**) Frequency and absolute numbers of SARS-CoV-2 Ag+ cells in the myeloid and epithelial compartments, determined by IMC, in the post-mortem lung samples according to HIV status within the COVID-19 group. **E**) Cell type enrichment analysis of the cell populations identified in Malawi lung IMC data. The comparison shown is between COVID-19 versus LRTD cases. To correct for multiple testing, the spatial false discovery rate (FDR) was calculated and only dots with spatialFDR < 0.05 are shown. **F**) Cellular landscape of histopathological lesions based on matched H&E and IMC analysis of post-mortem lung samples from the different clinical groups. The lesions were pooled and the graph shows the average proportion of each cell type in each lesion type. Colour key is as for Fig.2a,b.

**Extended data Figure 3 |** *Minimal SARS-CoV2 reads in single-cell data of lung, nasal and blood cells*

**(A)** Lung reference as in Fig 3a. **(B-D)** UMAPs indicate the cells in which we found reads that mapped to the SARS-CoV2 genome, coloured by case. **D)** Table showing absolute cell numbers per case that contain expression of valid UMI to the SARS-CoV2 genome in the lung, peripheral blood and nasal compartment.

**Extended data figure 4 |** *Lung cell proportions and gene module scores*

A-B) Cell type proportion bar plots of lung cell types in A) Immune cells and B) Stromal cells corresponding with Fig 3B and Fig 3C, grouped by disease group. C) Heatmap showing the mean gene module score across cells in gene sets associated with the *alpha, beta, gamma, lambda and TNF response*. Cell types have been grouped by COVID-19 and LRTD to show the difference in response and module score values have been scaled between -1 and 1.

**Extended data Figure 5** | *Heatmap of interferon response genes in lung*

Heatmaps showing the log fold change of up/down-regulated interferon response genes taken from immunologic gene sets involved in the immune response. Comparisons include the change in interferon response in cells from the HLCA COVID-19 cohort compared to the control cases **(A)**, our COVID-19 cohort compared to control cases from the HLCA **(B)** and interferon responses from our COVID-19 cohort compared to the HLCA COVID-19 cohort **(C)**.

**Extended data Figure 6** | *Bulk approaches to explore gene signatures*

Heatmaps showing cytokine signatures in different tissues. Values are plotted as z-score (grey mean not measured). Samples are grouped by their disease type. Luminex data of Nasal (**A**) and Plasma (**B**). Data were transformed with a log2 and for the visualistion with ComplexHeatmap in R with a Z-score by gene. For the statistical tests we compared levels of IFN-y, IL6, IL8, TNF and IL1b in nasal fluid and plasma between the Covid-19 and LTRD samples using a Welch Two Sample t-test which was non-significant for all comparisons, we did not correct for multiple comparisons. (**C-E)** Pseudobulk heatmaps showing cytokines included in the Luminex panel on the transcriptomic level in the peripheral blood, lung and nasal compartment per patient. As for Luminex we compared levels of IFN-y, IL6, IL8, TNF and IL1b in nasal, blood and lung cells between the Covid-19 and LTRD samples using a Welch Two Sample t-test which was non-significant for all comparisons, we did not correct for multiple comparisons.

**Extended data Fig 7** | *Cell-cell interactions in the IMC datasets*

**(A)** Heatmaps for the non-LRTD group showing co-localised cell types as shown by the IMC providing insight into potentially interacting cell types in the lung, shown for comparison with the same data from LRTD cases Fig.5b and Covid19 cases Fig5c (main figures) **(B-D)** Cellular maps showing the spatial location of specific immune cells and the potential of cell-cell interactions – highlighting spatially enriched macrophage interactions identified in Fig.5b (main figure). (**B**) shows interactions between alveolar macrophages (purple), apoptotic alveolar macrophages (blue) and apoptotic alveolar macrophages (yellow). C) shows interactions between apoptotic alveolar macrophages (yellow) fibroblast (lilach) and SARSCoV2+ Epithelial cells (purple) (**D**) shows interactions between activated endothelial cells (blude) and SARSCoV2+ Neutrophils (green)

**Extended data Table 1 |** *Characteristics of the patients*

Summary table of cases recruited into our study.

Abbreviations:

PMI = post mortem interval in hours

Obese/ Underweight indicates nutritional status, determined by a combination of abdominal circumference measurements and mid-arm circumference measurements and based on reference data for men and women in African populations : ↑ = overweight; ↑↑ = obese; ↑↑↑ = morbidly obese; ↓ = underweight; ↓↓ = severely underweight Pre-morbidity: DM2 = type II diabetes mellitus, HT = hypertension S.S. to death = symptom start to death, indicating the number of days between the first symptoms consistent with Covid19 (fever, cough, headache etc) and death. Lung IMC = Imaging Mass Cytometry, Lung sc = lung cell single-cell RNA-seq, nasal sc = Nasal cell single-cell RNA-seq, blood sc = blood cell single-cell RNA-seq, Nasal Lx – Nasal Luminex, is for Multiplexed cytokine array on nasal fluid. A dot for each of these parameters indicates that data are available for that assay for that case.

**Extended data Table 2** | *Comparison of cell proportions in IMC data*

The table is divided into three sections. The top section shows the proportion of different immune, stromal and vascular cells in the three groups in the Malawi cohort: Covid19, LRTD and non-LRTD and statistical comparison between the groups. The middle section shows comparison between HIV positive and HIV negative Covid19 cases. The bottom section shows comparison between the three IMC cohorts after integration: Brazil, USA and Malawi.

**Extended data Table 3** | *Differential gene expression in single-cell data by cell type* Differential gene expression analysis results from all cell types in the lung, nasal and blood tissue compartments. The table is organised with each comparison of cell types in the Malawi cohort in Covid-19 compared to LRTD in the three tissues and includes comparisons between lung cells in the Malawi cohort compared to the HLCA cohort. The table contains the average log fold change (avg_log2FC) along with the p-value (p_val) and multiple-test corrected p-values (p_val_adj).

## Notes

### Competing Interest Statement

The authors have declared no competing interest.

### Summary of Updates

Author affliations for some of the authors have been updated.

