## Extended figures for "Spatially resolved single-cell atlas of the lung in fatal Covid19 in an African population reveals a distinct cellular signature and an interferon gamma dominated response"

Extended data Fig. 1

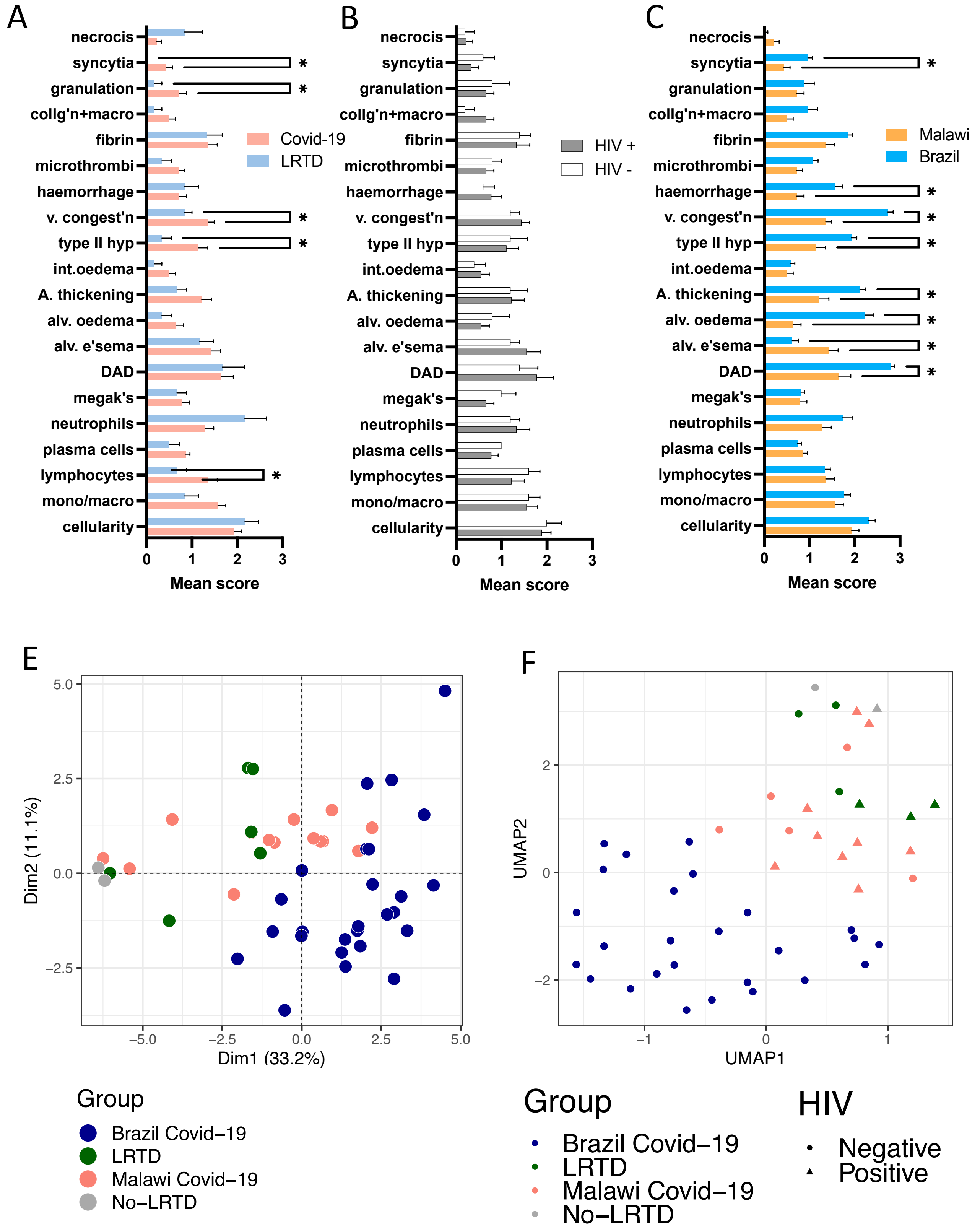

**Extended data figure 2:**

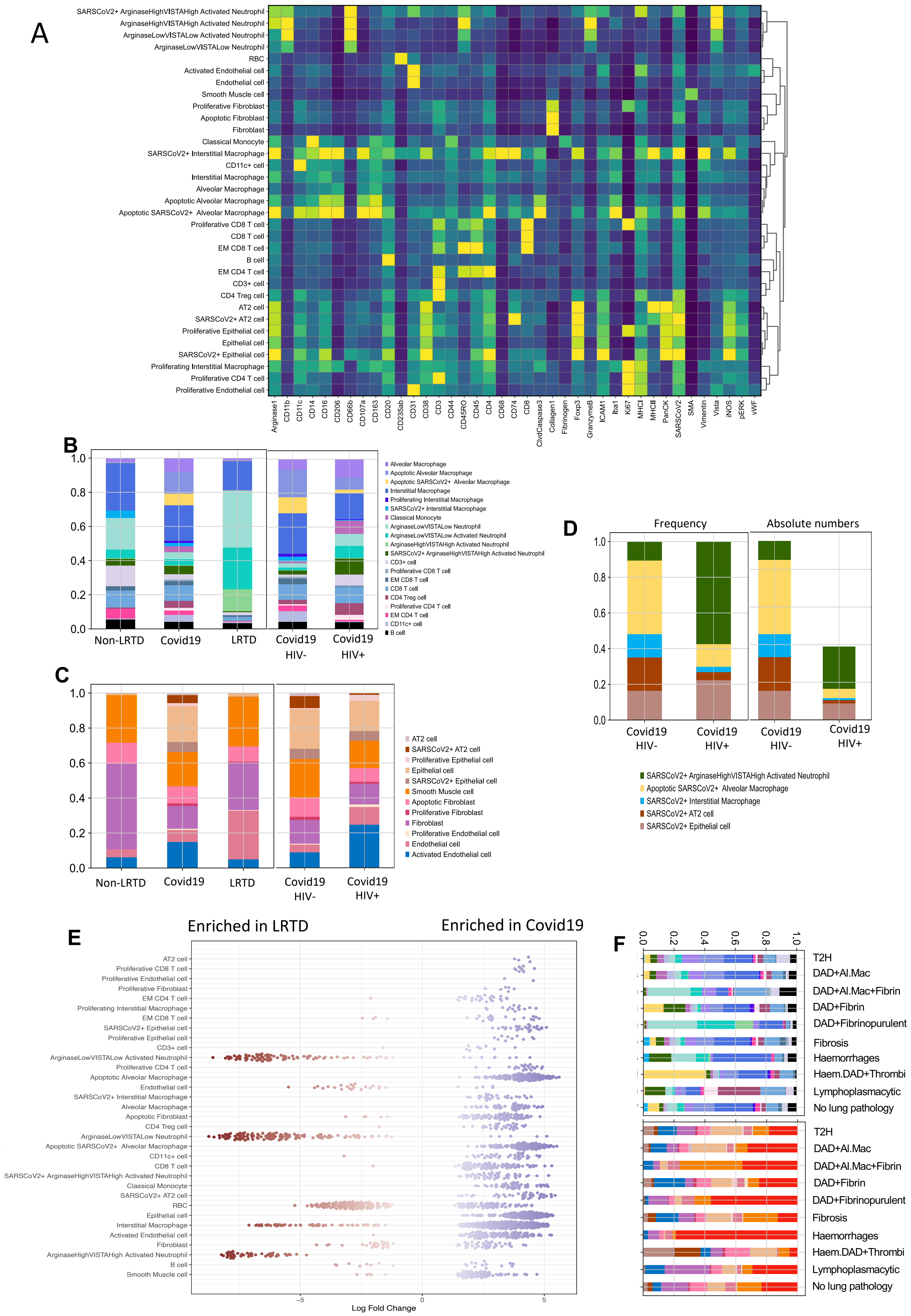

**Extended data figure 3:**

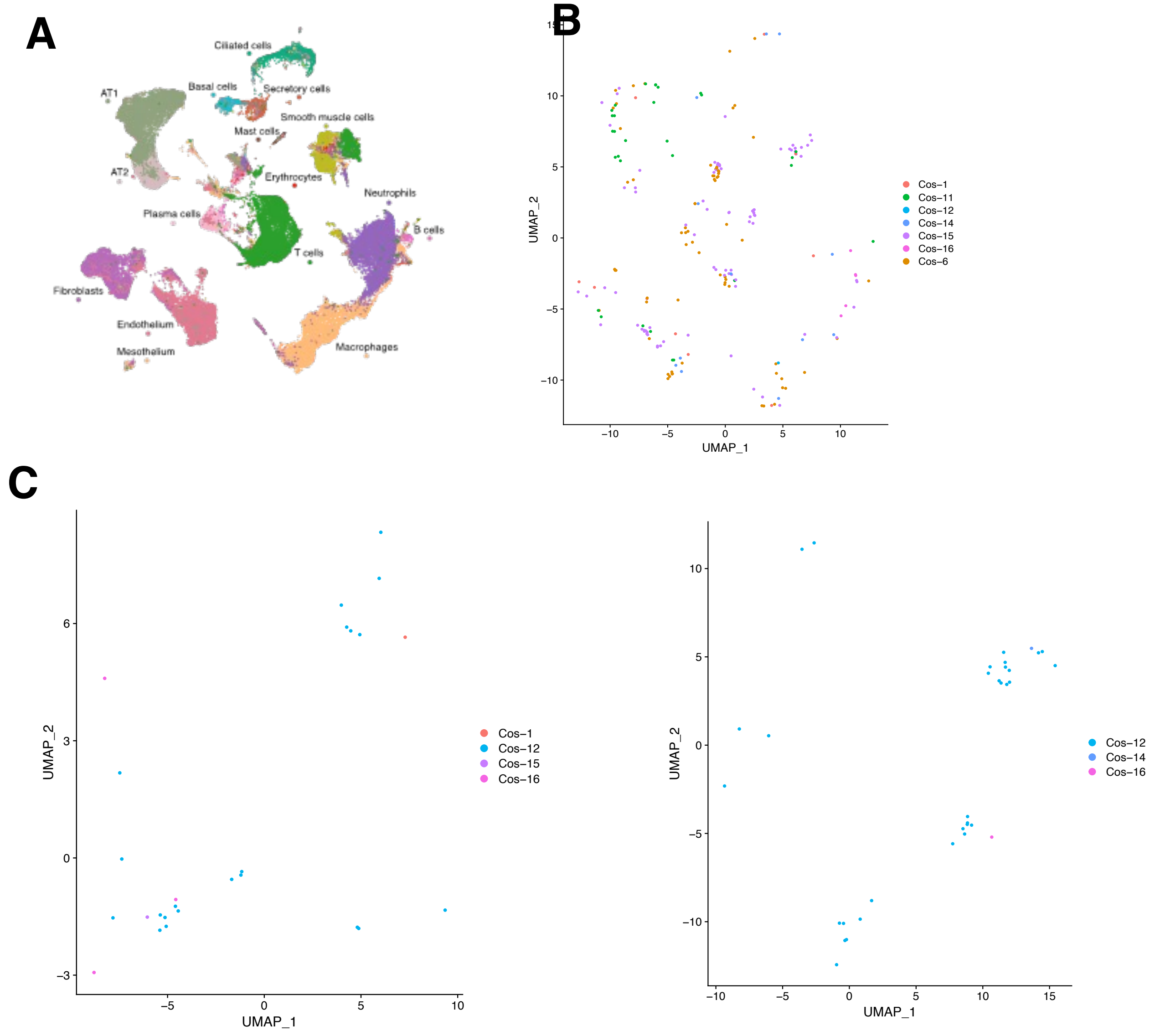


**D**

| Tissue | Lung | Blood | Nasal | Total |
|---|---|---|---|---|
| Cos1 | 10 | 1 | X | 11 |
| Cos6 | 69 | X | X | 69 |
| Cos11 | 30 | X | X | 30 |
| Cos12 | 1 | 21 | 32 | 54 |
| Cos14 | 12 | X | 1 | 13 |
| Cos15 | 87 | 1 | X | 88 |
| Cos16 | 5 | 3 | 1 | 9 |
| Total | 214 | 26 | 34 | 274 |

**Extended data Fig 4:**

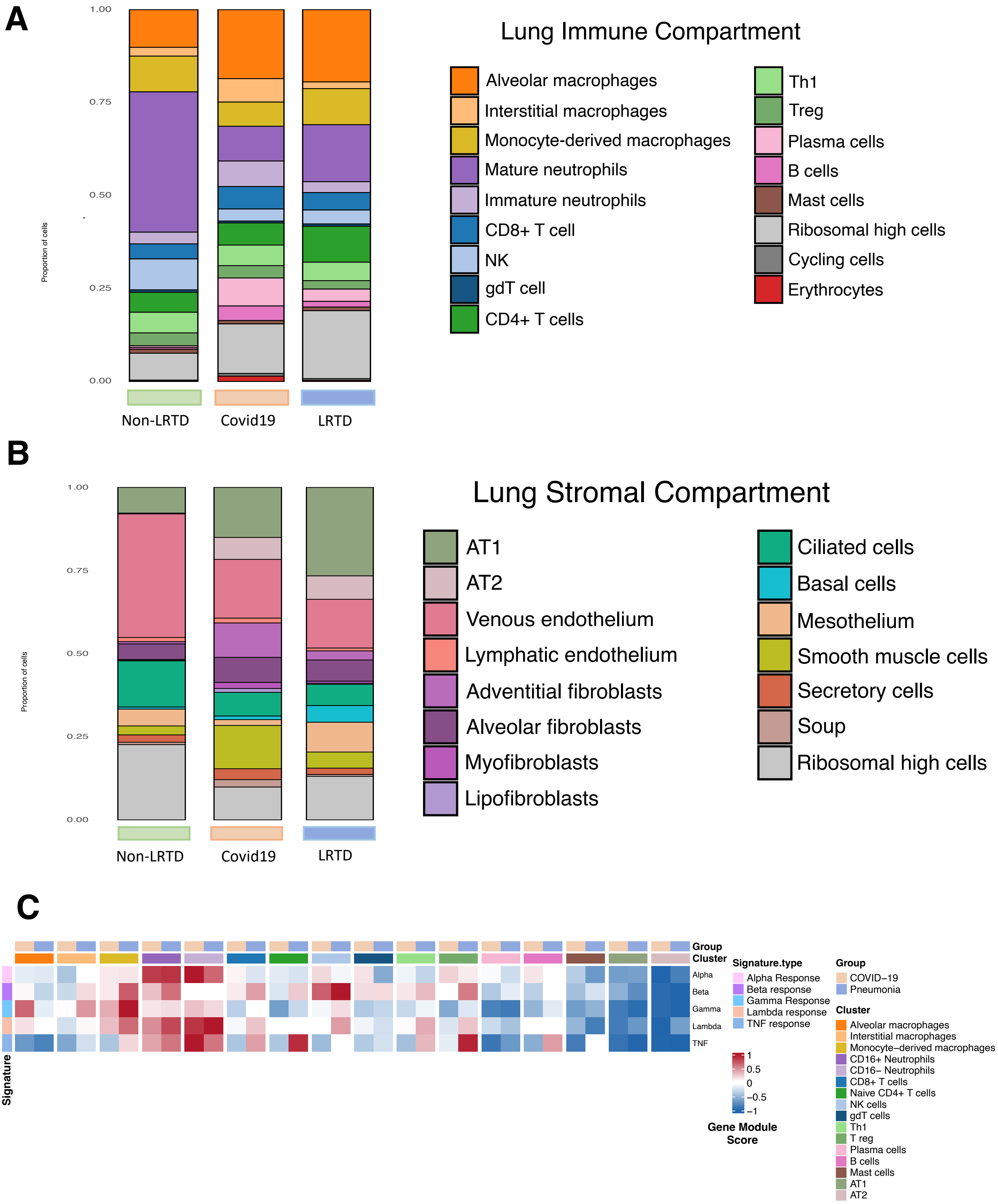

**Extended data Fig 5:**

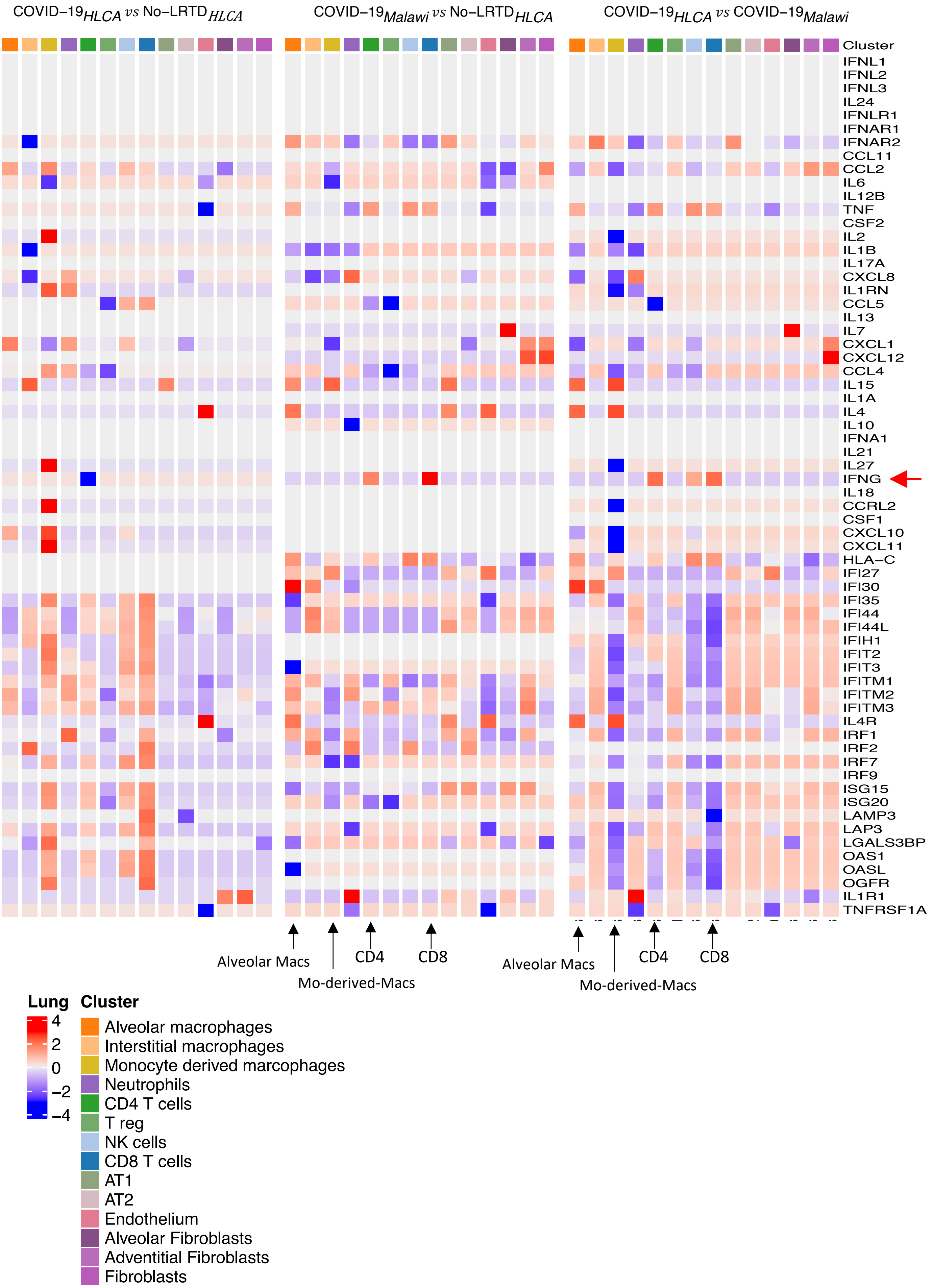

**Extended data Fig 6:**

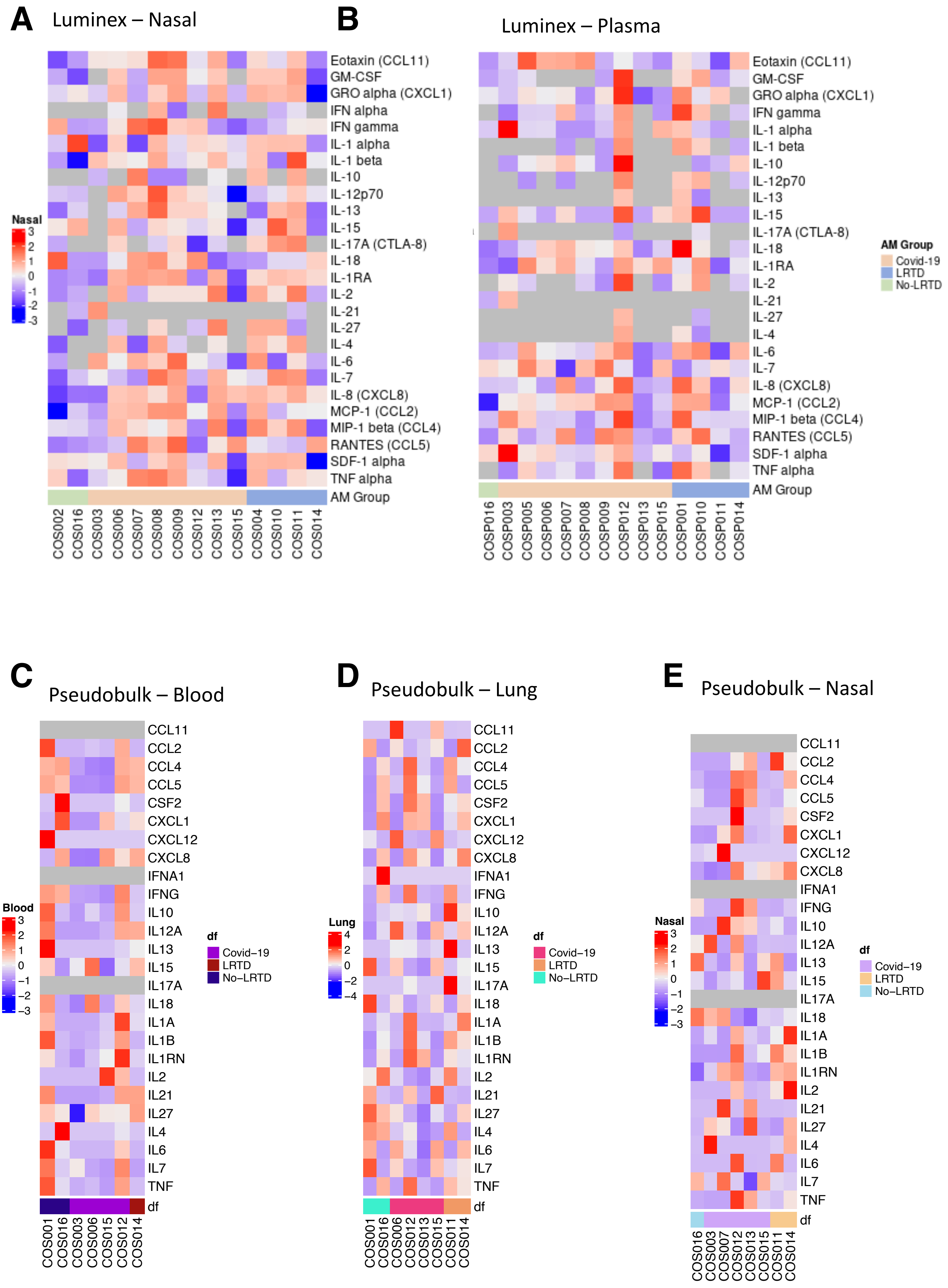

### Extended data Fig. 7

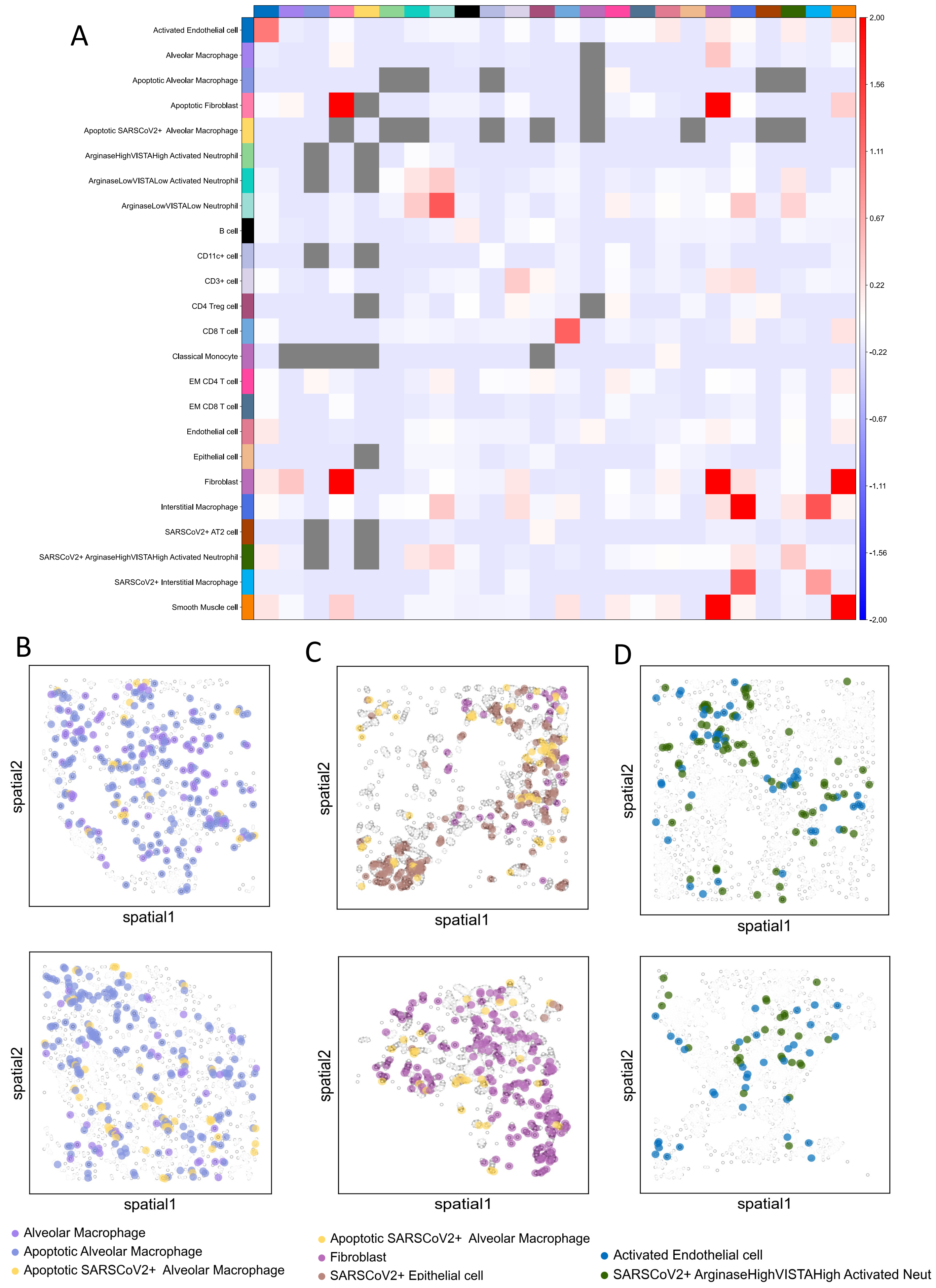

### Extended data Table 1

| | Case | Diagnosis | HIV | Sex | Age (yr) | PMI (hr) | Obese/Maln. | Pre-morbidity | S.S to death | Lung sc/sn | Nasal sc | Blood sc | Lung IMC | Nasal Lx | Blood Lx |
|---|---|---|---|---|---|---|---|---|---|---|---|---|---|---|---|
| **Covid-19** | **3** | C19 | 1 | M | 55-60 | 9 | ↑ | DM2, HT | 7 | | ● | ● | ● | ● | ● |
| | **5** | C19 | 1 | M | 55-61 | 5 | ↑↑ | DM2, HT | 6 | | | | ● | | ● |
| | **6** | C19 | 1 | M | 50-55 | 4 | ↑↑ | DM2, HT | 7 | ● | | ● | ● | ● | ● |
| | **7** | C19 | 1 | F | 50-55 | 5 | ↑↑↑ | Cancer | 4 | | ● | | ● | ● | ● |
| | **8** | C19 | 1 | F | 45-50 | 6 | ↑↑↑ | HT, A | 8 | | | | ● | ● | ● |
| | **9** | C19 | 0 | F | 50-55 | 5.5 | ↑↑ | None | 8 | | | | ● | ● | ● |
| | **12** | Sepsis+C19 | 0 | F | 60-65 | 9 | ↓ | None | 29 (2)* | ● | ● | ● | ● | ● | ● |
| | **13** | C19 | 0 | F | 70-75 | 5.5 | ↑↑ | DM2, HT | 20 | ● | ● | | ● | ● | ● |
| | **15** | C19 | 0 | M | 55-60 | 10.5 | → | None | 5 | ● | ● | ● | ● | ● | ● |
| **No LRTD** | **2** | TB | 1 | M | 45-50 | 9 | ↓↓ | None | 13 | ● | | ● | | | ● |
| | **16** | B. Pneum. | 0 | F | 60-65 | 2.5 | → | HT | 9 | | | | ● | ● | |
| **LRTD** | **1** | TB | 1 | F | 50-55 | 3 | → | None | 17 | | | | | ● | ● |
| | **4** | L. Cancer | 1 | F | 60-65 | 3 | ↓↓↓ | None | 10 | ● | ● | | ● | ● | ● |
| | **10** | B. Pneum. | 0 | M | 60-65 | 9.5 | ↓ | HT | 5 | ● | ● | ● | ● | ● | ● |
| | **11** | Sepsis | 1 | F | 50-55 | 10.5 | ↓↓ | None | 4 | | | | ● | ● | |
| | **14** | Stroke | 0 | F | 50-55 | 9 | → | None | 2 | ● | ● | ● | ● | ● | ● |
